# Computer Designed PRC2 Inhibitor, EBdCas9, Reveals Functional TATA boxes in Distal Promoter Regions

**DOI:** 10.1101/2020.11.16.385922

**Authors:** Shiri Levy, Logesh Somasundaram, Infencia Xavier Raj, Diego Ic-Mex, Sven Schmidt, Ammar Alghadeer, Henrik Honkanen, R. David Hawkins, Julie Mathieu, Yuliang Wang, David Baker, Karol Bomsztyk, Hannele Ruohola-Baker

## Abstract

The critical process in development, bifurcation of cellular fates, requires epigenetic H3K27me3 marks propagated by PRC2 complex. However, the precise chromatin loci of functional H3K27me3 marks are not yet known. Here we identify critical PRC2 functional sites at a single nucleosome resolution. We fused a computationally designed protein, EED binder (EB) that competes with EZH2 and thereby disrupts PRC2 function, to dCas9 (EBdCas9) to direct PRC2 inhibition at a precise locus using gRNA. We targeted EBdCas9 to 4 different genes (*TBX18, p16, CDX2* and *GATA3*) and observed epigenetic remodeling at a single nucleosome resolution resulting in gene activation. Remarkably, while traditional TATA box is located 30bp upstream of TSS, we identified a functional TATA box, >500bp of TSS, normally repressed by PRC2 complex. Deletion of this TATA box eliminates EBdCas9 dependent TBP recruitment and transcriptional activation. Targeting EBdCas9 to *CDX2* and *GATA3* results in trophoblast trans-differentiation. EBdCas9 technology is broadly applicable for epigenetic regulation at a single locus to control gene expression.

**Graphical Abstract:** 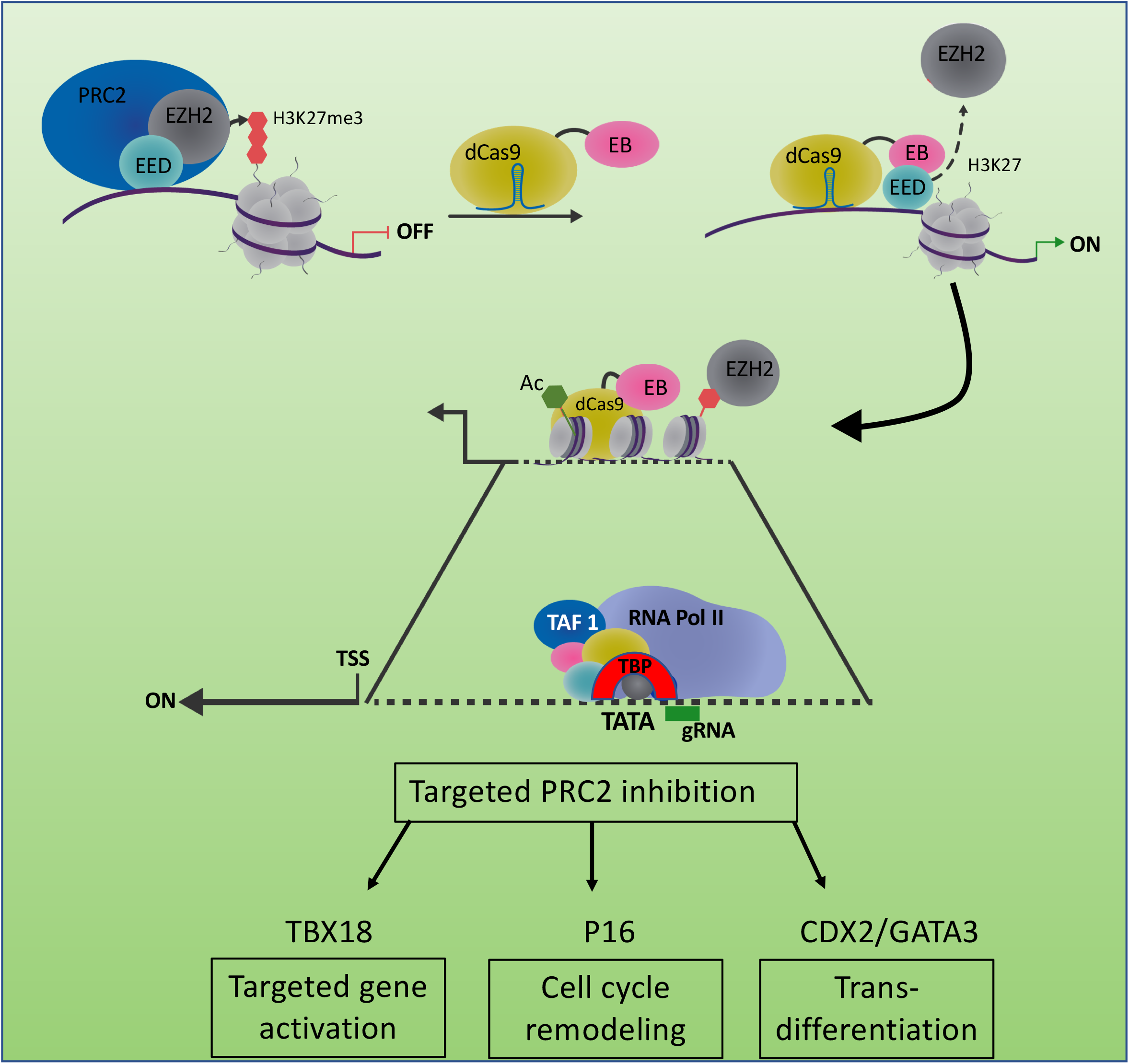

## Introduction

A central question in epigenetics and developmental biology is the role of specific histone 3 lysine 27 methylation (H3K27me3) marks in cell fate decisions^1^. Polycomb repressive complex 2 (PRC2) is an evolutionarily conserved, repressive H3K27me3 methyltransferase complex that plays a key role in developmental transitions. While broad upstream regions of developmental genes are decorated with H3K27me3 marks it is not known which, if any, single nucleosomes require H3K27me3 marks for gene repression and cell fate determination.

The two main complexes involved in Polycomb based repression are PRC1 and PRC2. PRC1 catalyzes monoubiquitylation of histone H2A Lys 119 (H2AK119ub) while PRC2 catalyzes the mono-, di-, and trimethylation of histone H3 Lys27 (H3K27me1/me2/me3) ^2^. Recent structural analysis has revealed the mechanism of PRC1/PRC2 spatiotemporal regulation and H3K27me3 mark spreading^3, 4^. Non-canonical PRC1 containing RYBP ubiquitinates H2K119 to recruit accessory proteins JARID2 and MTF2 ^5–7^. JARID2 recruits PRC2 and contributes to allosteric activation of PRC2 ^8^. PRC2 containing EED and EZH2 facilitates H3K27me3 marks by reading (EED) and writing/catalyzing (EZH2) new marks. H3K27me3 marks are then read by canonical PRC1 containing CBX, resulting in chromatin compaction, H2AK119ub and gene repression ^9–15^. During developmental transitions the chromatin undergoes extensive epigenetic remodeling at promoter and enhancer regions, where H3K27me3 marks are added or removed to produce the required increase or decrease in gene expression ^16, 17^. Importantly, only the repressive marks, but not the activating histone tail marks, are instrumental for epigenetic inheritance during development ^18^. Recent studies suggest that proximal regulatory marks governing gene expression are associated with histone modification domains within 2.5 kb of known transcriptional start sites of RefSeq genes ^19^. However, it is not known if any specific H3K27me3 marked nucleosomes are critical for control of gene expression or if the entire broad 2.5kb region is required for gene repression. This issue has been challenging to address as existing genetic and chemical methods eliminate all H3K27me3 marks, without precision.

Recent studies that fuse deadCas9 (dCas9) to proteins that promote or remove histone methyl groups provide tools for manipulating the epigenome. For example, the repressor KRAB recruits methyltransferases to increase H3K9me3 repression levels ^20, 21^, and fusing KRAB to dCas9 with the appropriate guide RNA (gRNA) results in 50%-99% gene repression in various cell lines, primary cultures, and pluripotent stem cells^22–27^. Fusion of dCas9 to the transcriptional activators VP16, VP64, and VP65 promotes the recruitment of chromatin modifiers, resulting in chromatin decondensation, accumulation of histone marks such as H3K27ac and H3K4me3; binding of Pol II; and subsequent mRNA transcription^28, 29^. Fusion of dCas9 to the histone demethylase LSD1/KDM1A responsible for histone K4 and K9 methyl group removal, affects ESC self-renewal and differentiation as well as cancer cell proliferation and development ^23, 30, 31^. Fusion of dCas9 to the PRC2 methyltransferase (EZH2) did not result in repressive activity unless co-targeted with DNMT3A-dCas9 and induced with DNMT3L for prolong period of time. This suggests that histone methylation alone is not sufficient to produce gene repression ^32, 33^. Interestingly, recent studies that fuse Cas9 to JARID2 alone resulted in enrichment of EZH2 and H3K27me3 at targeted loci and 20% reduction in gene expression ^34^.

Despite the considerable array of tools available to modify the epigenome, currently there is no method to inhibit PRC2 function at a particular genomic locus and precisely at a single nucleosome in order to determine which specific H3K27me3 marks result in transcriptional repression. We have generated a computer designed protein that binds EED (EED binder; EB) and thereby competes with EZH2 localized activity ^35^. By fusing the designed PRC2 inhibitor, EB, to dCas9, we enable probing H3K27me3 function at precise gene loci in the natural biological context of transcription repression. Here we show that EBdCas9/gRNA is able to affect the PRC2 complex, remodel the epigenetic marks of specific target sites, upregulate genes of interest, and promote epigenetic memory. We reveal precise loci for PRC2 requirement for transcription repression. In the case of *TBX18* gene this precise PRC2 activity normally represses distal TATA box region. We also utilized EBdCas9 to explore the role of H3K27me3 modifications in epigenetic control of developmental cell fate bifurcation to inner cell mass (ICM) or trophectoderm (TE) ^36, 37^. Significantly, we now identify the precise location of H3K27me3 marks critical for TE trans-differentiation from iPSCs using the targeted PRC2 inhibitor EBdCas9.

## Results

### EBdCas9/gRNA activates *TBX18* transcription

The catalytic and substrate recognition functions of PRC2, mediated by the SET domain containing EZH2 subunit and the tri-methyl lysine binding EED subunit, respectively, are coupled by binding of the N-terminal helix of EZH2 to an extended groove on EED^38^. Disrupting PRC2 function can be achieved by chemical drugs or knock out of key PRC2 components ^39, 40^. We previously generated and characterized a computationally designed protein (EED Binder; EB) that binds to the EZH2 binding site on EED by two orders of magnitude higher affinity than EZH2^35^. As a control, we created an EED binder negative control (NC), where two amino acid mutations: F47E and I54E on the EED binding interface abolish binding to EED^35^. EB, but not NC, with subnanomolar affinity, forms tight complexes with EED, reduces EZH2, and JARID2 global levels, and exhibits a significant genome wide reduction of H3K27me3 repressive marks in promoter regions ^35^.

To target EB to specific chromatin locus to test its functionality in precise regions, we fused EB to dCas9 (EBdCas9). EBdCas9 together with targeted guide RNA (gRNA) now allows us, for the first time, to disrupt PRC2 at a local level and identify which precise regions within H3K27me3 domains, if any, are required for controlling gene expression of the targeted gene of interest (**Fig 1A**). To generate EBdCas9 protein, we fused EB to dCas9 under AAVS1-TREG inducible promoter of dCas9-NLS-mCherry plasmid ^41^ (**Fig 1B**). Similarly, we fused the NC control to dCas9 to critically distinguish between dCas9 non-specific effects on chromatin and EB specific effects on PRC2. A 30 aa residue 6×5 (SGGGG) linker was inserted between EB or NC and dCas9 for free mobility and permissive binding action once EB/NCdCas9-gRNA is bound to targeted DNA. The EB-linker-dCas9-NLS-mCherry (EBdCas9) and NC-linker-dCas9-NLS-mCherry (NCdCas9) constructs were transformed to iPSC (WTC) using TALENS to enforce recombinant homology at the safe harbor locus, AAVS1 site on chromosome 19. Following antibiotic selection, the lines were validated for EBdCas9 and NCdCas9 mCherry expression with or without Dox induction where no leakiness was observed and stem cell morphology was maintained (**Fig 1C**). Unlike EB, that causes global EZH2 and H3K27me3 reduction in hESC^35^, WTC cells are not affected by EBdCas9 when induced without gRNA, and EZH2 and H3K27me3 levels remain the same between induced and uninduced EBdCas9 (**Fig 1D**). EBdCas9 and NCdCas9 transcript expression was found to be 50x lower compared to EB or NC suggesting that construct based off-target effects may be minimal with the EBdCas9 and NCdCas9 constructs (**Sup 1A**). Since Cas9 is shown to undergo conformational changes upon binding to DNA^42^, it is also plausible that EBdCas9 fusion may comprise conformational steric hindrance effects that do not allow EB to bind promiscuously to EED and therefore no global H3K27me3 or EZH2 reduction is observed.

**Figure 1.**
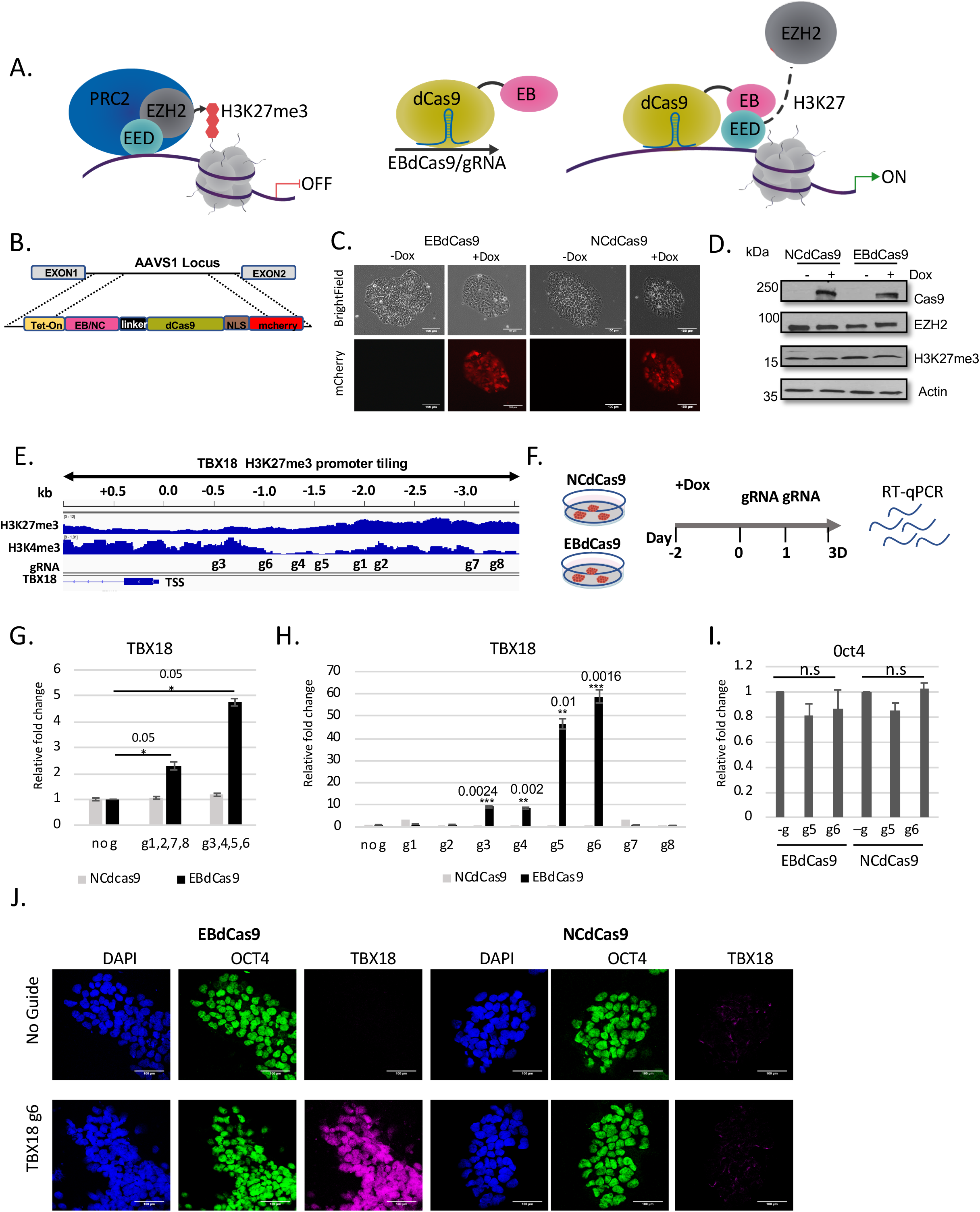
EBdCas9 in precise loci upregulates TBX18 expression. **A**. Model of EBdCas9 precise elimination of PRC2 activity in targeted loci **B**. EBdCas9-mCherry and NCdCas9-mCherry construct under Tet-On operator. **C**. Generation of stable EBdCas9 or NCdCas9 transgenic iPSC lines after 3D induction of (2μg/ml) Doxycycline (Dox). **D**. Immunoblot analysis of EBdCas9 and NCdCas9 whole cell lysate after 3D Dox induction. **E**. Integrative Genomic Viewer of *TBX18* H3K27me3 and H3K4me3 promoter tiling. **F**. Timeline of EBdCas9 or NCdCas9 induction and gRNA transfection. **G-I**. RT-qPCR analysis of TBX18 or Oct 4 expression for EBdCas9 and NCdCas9 normalized to beta-Actin and calculated as relative fold change compared to no guide (induced with Dox) of each respected cell line (**G**) after cocktail TBX18 gRNA transfection with either g1,2,7,8 or g3,4,5,6 TBX18 promoter tiling, (**H**) after individual TBX18 gRNA (1-8) transfection. (**I**) after individual transfection of TBX18 gRNA g5 and g6. \**p*<0.05, ** *p*<0.01, \*\*\**p*<0.001 one-way ANOVA performed. n=3 biological replicates. **J**. Immunofluorescent imaging of EBdCas9 WTC and NCdCas9 WTC for either no guide or after transfection with TBX18 gRNA 6 (TBX18 g6). Blue-Dapi, Green-Oct4, Far red- TBX18; scale bar is 100*μ*m.

To identify the effects of targeting EBdCas9 or NCdCas9 to specific chromosomal loci we screened for genes that showed the most H3K27me3 reduction after EB treatment previously tested in ChIPseq H3K27me3 EB analysis ^35^. *TBX18*, a growth promoting transcription factor of the sinoatrial node T-box 18, is required for embryonic development and conversion of working myocytes into sinoatrial cells ^43, 44^. *TBX18* was observed as a highly significant upregulated gene with reduced H3K27me3 marks after EB expression and was therefore selected as a candidate locus to analyze the action of EBdCas9 construct (**Sup 1B**) ^35^. In iPSCs, the *TBX18* gene promoter shows bivalency- the gene upstream region is simultaneously decorated with both H3K27me3 repressive marks and H3K4me3 active marks. We tiled the *TBX18* upstream promoter region, based on UCSC Genome Browser (GRCh36/hg36) Assembly, with guides chosen to be specific to the region using CRISPRscan^45^ gRNAs prediction tool. Chosen guides were used with EBdcas9 and NCdCas9 to screen for loci sensitive for PRC2 targeted-locus-activation (**Fig 1E**). EBdCas9 was induced at day -2 using doxycycline and transiently transfected with *in vitro* synthesized gRNA at day 0 and day 1 and the cells were collected at day 3 (**Fig 1F**). To initiate the screens to test which upstream regions are more sensitive to EB action and therefore more likely to activate transcription, we transfected the cells with a combination of guides. Interestingly, while guides 1.9-3.5kb upstream from TSS (g1,2,7,8,) did not show significant effects in *TBX18* transcription, the guides 0-1.9kb from TSS (g3,4,5,6) showed 4 fold *TBX18* upregulation compared to no guide treatment (**Fig 1G**). The control NCdCas9 did not show effects with either groups of guides (**Fig 1G**).

To dissect the EB responsive region in a more precise manner, we transfected the cells with higher levels of each guide individually and analyzed *TBX18* transcriptional increase. Induction of EBdCas9 with the individual *TBX18* gRNAs resulted in *TBX18* transcript upregulation between 10 fold (gRNA 3 and 4) and 50-60 fold (gRNA 5 and 6) compared to no guide, EBdCas9 (g1,2,7,8), NCdCas9 or KRABdCas9 **(Fig 1H; Sup 1C**). gRNAs 3-6 (−0.5kb to-1.5kb) localized to unique chromatin domain where H3K4me3 marks are low, while H3K27me3 marks and EZH2 are still enriched (**Fig 1E, Sup 1D**). To ensure all tiled gRNAs are equally accessible to targeted *TBX18* DNA, we used hESC Elf iCas9 cell line^46^ to transiently transfected different gRNAs and to analyze DNA accessibility by sequencing based InDel analysis at targeted site. To test the specificity of EBdCas9, we monitored *POU5F1*(OCT4) gene expression and observed no significant changes in *OCT4* in cells treated with gRNA targeting *TBX18* promoter (**Fig 1I**), hence *TBX18* upregulation is not a secondary effect of cellular differentiation. Accordingly, TBX18 protein over expression was detected with EBdCas9/g6, but not with NCdCas9/g6 (**Fig 1J**). ATACseq analysis of EBdCas9/g6 or NCdCas9 /g6 transfected cells resulted in similar open chromatin region. Therefore, since no significant change in chromatin accessibility were monitored we concluded that that transcript upregulation is dependent on PRC2 local changes (**Sup 1B**). We conclude that EBdCas9, but not NCdCas9 is able to activate *TBX18* gene expression at precise loci.

### EBdCas9/gRNA remodels epigenetic marks and epigenetic memory in TBX18 locus

To dissect the mechanism of EBdCas9 action at precise genomic locus, we used ChIP-qPCR (PIXUL-Matrix-ChIP ^47^) to analyze the epigenetic landscape of *TBX18* g6 targeted region. As above, EBdCas9 or NCdCas9 expression was induced in iPSC (WTC) using doxycycline followed by *TBX18* gRNA6 (g6) transfections at day 0 and day 1 and analysis on day 3 by RT-qPCR and ChIP-qPCR (**Fig 2A**). As expected, while both constructs showed similar dCas9 expression, only EBdCas9/g6 showed significant increase of *Tbx18* transcription (**Fig 2B**). Using ChIP-qPCR method, we quantified the abundance of EB/NCdCas9 (Cas9), Histone3 with K27me3 marks (H3K27me3) and EZH2 proteins at TBX18 g6 region by generating and quantifying the 150bp amplicon surrounding g6 chromatin region. The ChIP-qPCR assay using Cas9 antibody (Ab) confirms that both EBdCas9 and NCdCas9 are recruited to guide 6 locus (**Fig 2C**). However, only EBdCas9 results in reduction of EZH2 at g6 locus (**Fig 2C**), most likely by directly blocking EZH2 binding to EED^35^. This resulted in reduction of H3K27me3 marks in EBdCas9/g6 at guide 6 specific locus (**Fig 2C**). These data show that EBdCas9 is able to disrupt EED-EZH2 interaction at precise locus.

**Figure 2.**
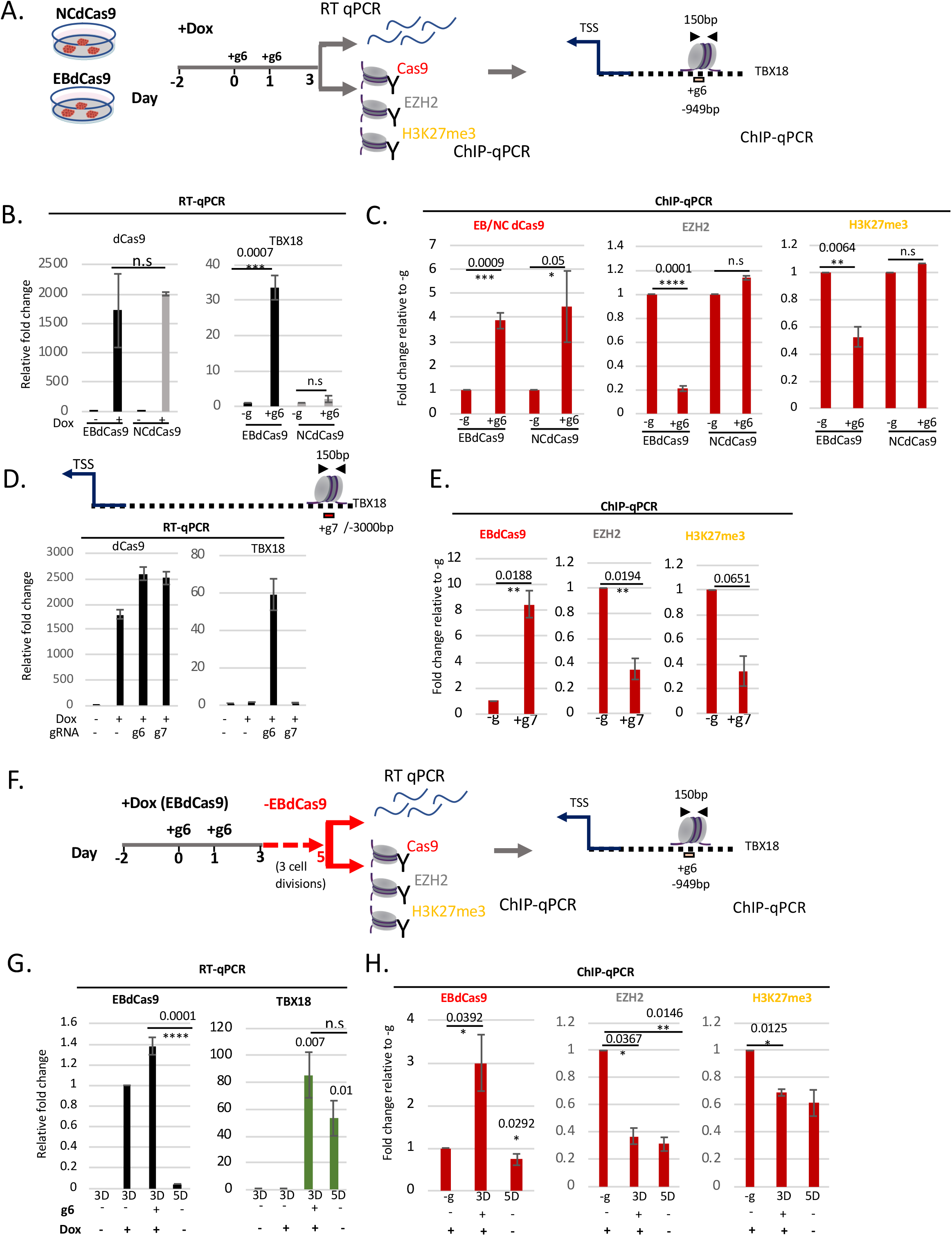
EBdCas9 eliminates EZH2 in PRC2 complex, remodels epigenetic marks and actuates epigenetic memory on TBX18 locus. **A**. EBdCas9 and NCdCas9 timeline for Dox induction, gRNA transfection and analysis: RT-qPCR or ChIPqPCR using the antibodies Cas9, EZH2 and H3K27me3 and analyzing TBX18 g6 DNA region ~1.0kb upstream of TSS (150bp). **B.**RT-qPCR of dCas9 or TBX18 relative fold change after 3D Dox induction (dCas9) or 3D TBX18 g6 RNA transfection (TBX18) normalized to beta-Actin and compared to no guide (Dox induced for TBX18) of each respected cell line. **C**. ChIP-qPCR of induced (+Dox) EBdCas9 and NCdCas9 after 3D transfection with TBX18 g6 RNA (+g6) or no transfection (-g). Normalized to fractioned Input /H3 and compared to −g relative fold change. Antibodies that were used for ChIP are listed above the graphs (Cas9, EZH2, H3K27me3) and the genomic region analyzed by qPCR includes TBX18 g6 locus. \**p*<0.05, ** *p*<0.01, \*\*\**p*<0.001 one-way ANOVA performed. n=3 biological replicates. **D.** RT-qPCR of dCas9 and TBX18 relative fold change after 3D Dox induction and TBX18 g6 RNA (+g6) or g7 RNA (+g7) transfection normalized to beta-Actin and compared to no guide (Dox induced) (-g) exactly as described in A. **E**. ChIP-qPCR of induced (+Dox) EBdCas9 after 3D transfection with TBX18 g7 RNA (+g7) or no transfection (-g). Normalized to fractioned Input /H3 and compared to −g relative fold change. Antibodies that were used for ChIP are listed above the graphs (Cas9, EZH2, H3K27me3) and the genomic region analyzed by qPCR includes TBX18 g6 locus. \**p*<0.05, ** *p*<0.01, \*\*\**p*<0.001 one-way ANOVA performed. n=3 biological replicates. **F**. EBdCas9 timeline for measuring epigenetic memory. EBdCas9 induction and TBX18 g6 RNA transfection exactly as in A. On Day 3 EBdCas9 media is replaced with no Dox (-EBdCas9) for 2 days and analyzed on Day 5 for RT-qPCR and ChIP-qPCR **G**. RT-qPCR analysis of EBdCas9 and TBX18 for 3 days (3D) or 5 days (5D) while inducing with Dox (+) or not (−) and in the presence of TBX18 g6 RNA (g6) (+) or not (−). **H**. ChIP-qPCR of no guide (-g), 3 days (3D) or 5 days (5D) EBdCas9 either induced with Dox (+) or not (−) or transfected with TBX18 g6 RNA (+) or not (−). Normalized to fractioned Input /H3 and compared to −g relative fold change. Antibodies that were used for ChIP are listed above the graphs (Cas9, EZH2, H3K27me3) and the genomic region analyzed by qPCR includes TBX18 g6 locus. \**p*<0.05, ** *p*<0.01, \*\*\**p*<0.001 one-way ANOVA performed. n=3 biological replicates.

To test whether EBdCas9 has access to chromatin in loci where transcription is not affected, we targeted EBdCas9 to TBX18 guide 7(g7) region (**Fig 1H, Fig 2D**). ChIP-qPCR assay of EBdCas9/g7 confirmed recruitment of Cas9, reduction of EZH2 and reduction of H3K27me3 marks at g7 locus (**Fig 2E, Sup 2A)**. These data show that while EBdCas9 is able to compromise PRC2 complex in g7 region, the PRC2 activity in this precise locus is not essential for transcription inhibition.

Recent data have shown that PRC2 dependent marks are faithfully inherited during cell division^18^. To learn whether EBdCas9 dependent reduction of EZH2 occupancy and H3K27me3 marks are inherited from the cell from which they descended (epigenetic memory), we repeated the assay as before, but allowed the cells to grow for additional 2 days after (3 cell cycles; **Sup 2B**) after eliminating EBdCas9/gRNA expression. The newly replicated cells were harvested at day 5 for RT-qPCR and ChIP-qPCR analysis (**Fig 2F**). RT-qPCR shows that EBdCas9 transcript is upregulated at day 3 post transfection (3D) but not detected 2 days after elimination of induction (5D) (**Fig 2G**). Using PIXUL-ChIP we show that while EBdCas9 protein is enriched in g6 region at 3D, no EBdCas9 enrichment is detected at 5D timepoint. We therefore proceeded to analyze the *TBX18* chromatin locus and transcript in these conditions. *TBX18* transcript showed 80 fold increase at 3D, furthermore, in contrast to EBdCas9, *TBX18* was still significantly upregulated at 5D (50 fold increase) (**Fig 2G**). To test if *TBX18* transcript detection 2 days after EBdCas9 elimination correlated with epigenetic memory, we performed PIXUL-ChIP for EZH2 protein and H3K27me3 marks at g6 region at 3D and 5D. ChIP-qPCR assay showed a depletion of both H3K27me3 and EZH2 at 3D, as expected. Importantly, this reduction of EZH2 and H3K27me3 persisted until 5D (**Fig 2H**), showing an epigenetic memory of PRC2 elimination on g6 region. These data reveal that EBdCas9 not only remodels the epigenome but also leads to epigenetic memory.

### EBdCas9 causes epigenetic spreading

To learn whether (i) EBdCas9/TBX18g6 epigenetic changes are limited to guide 6 chromatin locus, or whether (ii) the epigenetic changes spread to the neighboring regions, and whether (iii) the spread chromatin changes are inherited upon cell division, we tiled *TBX18* genomic area with primer sets between TSS and g6, as well as upstream from g6. As expected, at 3D, but not at 5D (2 days after eliminating EBdCas9 induction) EBdCas9 (Cas9) is localized to the g6 region (**Fig 3A**). In neither timepoint was EBdCas9 observed in any other loci upstream of *TBX18* TSS. However, EZH2 protein was significantly downregulated not only at g6 but also in loci between TSS and g6 both at 3D and 5D timepoints (**Fig 3B**). Since EBdCas9 was specifically localized to g6 region at 3D, but not at 5D, EZH2 reduction in these regions suggest that PRC2 disruption by EBdCas9 at g6 at 3D (i) spreads towards TSS and (ii) is inherited during cell divisions. Accordingly, disruption of the PRC2 complex also resulted in ongoing reduction of H3K27me3 marks between TSS and g6 (**Fig 3C**). An important PRC2 component, JARID2, showed reduction binding to chromatin between TSS and g6 at 3D timepoint (**Fig 3D**). Since JARID2 is critical for PRC2 homing and stimulation of EZH2 catalytic activity, its reduction in this region is further indication for reduced PRC2 activity ^48^. EED on the other hand, binds to EBdCas9 and remains at guide 6 at 3D, however it is depleted between TSS and g6 (**Fig 3D**). These data suggest that PRC2 disruption and thereby reduction of H3K27me3 epigenetic marks both in 3D and 5D timepoints are spreading from g6 towards the TSS.

**Figure 3.**
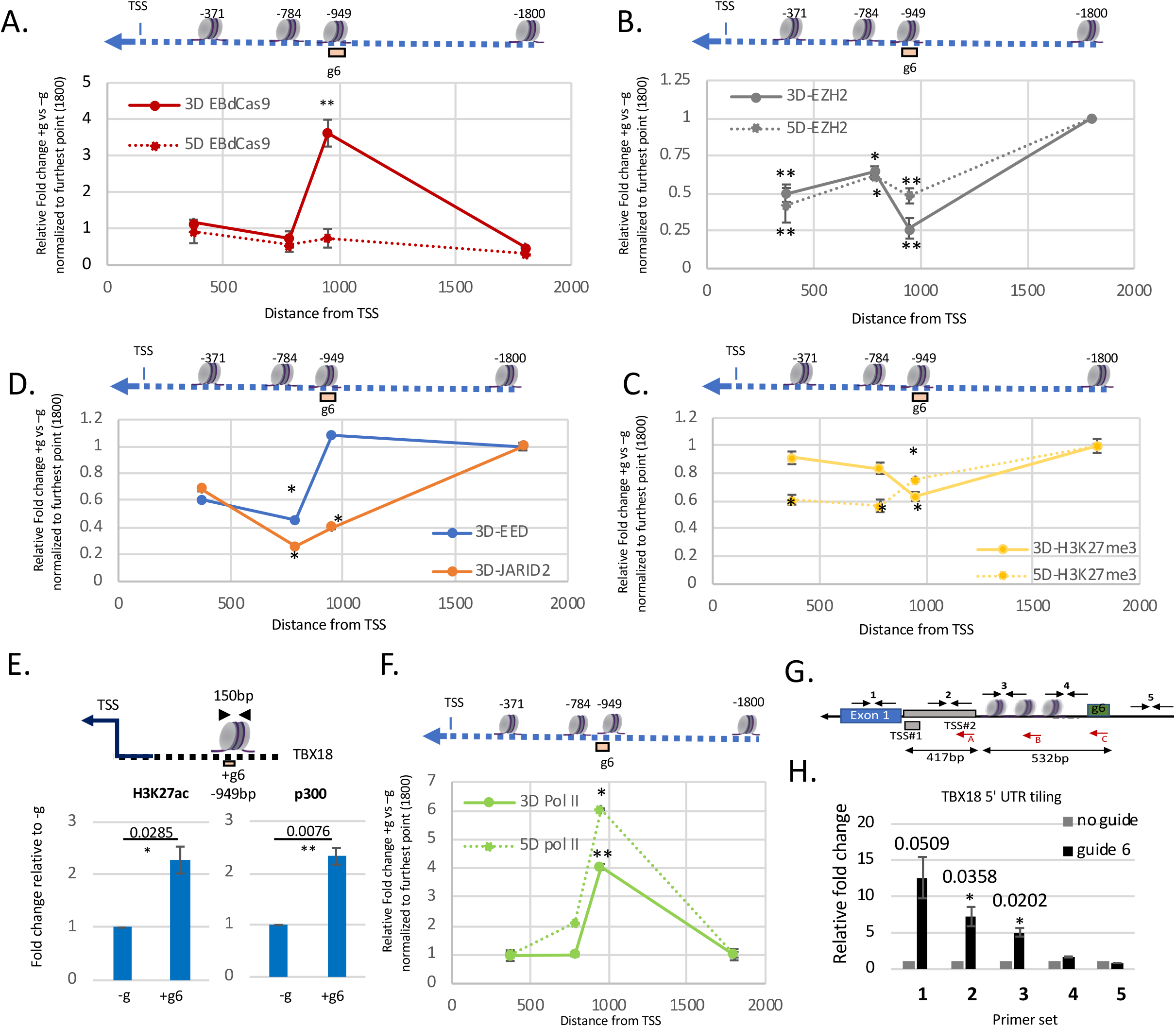
EBdCas9 causes epigenetic spreading and reveals distal TATA box function. **A-D** Tiling of EBdCas9/+g6 ChIP (**A**) Cas9, (**B**) EZH2, (**C**)H3K27me3, (**D**) EED and JARID 2 on TBX18 genomic loci (bp) relative to TSS (listed above nucleosome) using qPCR; solid red lines are 3D and dash red lines are 5D. Each point is relative fold change TBX18 g6RNA (+g6) vs no guide (-g), normalized to respected fractioned Input/H3 and compared to relative fold change using −1800bp primer set control. \**p*<0.05, \*\**p*<0.01, \*\*\**p*<0.001 one-way ANOVA performed. n=3 biological replicates. **E**. ChIP-qPCR of 3D induced EBdCas9 with TBX18 g6 RNA (+g6) transfected or not (-g). Normalized fractioned to Input/H3 and compared to no guide (-g). Antibodies that used for ChIP are listed above the graphs (H3K27ac and p300) and the genomic region analyzed by qPCR is TBX18 g6 locus. \**p*<0.05, \*\**p*<0.01, \*\*\**p*<0.001 one-way ANOVA performed. n=3 biological replicates. **F**. Tiling of EBdCas9/+g6 ChIP (Pol II CTD) on TBX18 genomic loci(bp) relative to TSS (listed above nucleosome) using qPCR exactly as in A; solid green lines are 3D and dash green lines are 5D (as described in 3A-D). Each point is relative fold change TBX18 g6RNA (+g6) vs no guide (-g), normalized to respected fractioned Input/ H3 and compared to relative fold change using −1800bp primer set control. \**p*<0.05, \*\**p*<0.01, \*\*\**p*<0.001 one-way ANOVA performed. n=3 biological replicates. **G**. Illustration of TBX18 TSS peak position relative to guide 6 (g6) region in bp. A, B, and C red arrows denote the reversed primer used for cDNA reverse transcription. Internal black arrows with Arabic numbers denote the PCR amplicon generated following cDNA production: C corresponds to primer sets #4; B corresponds to #3; A corresponds to primer sets #2,#1 and #5. **H**. RNA expression of *TBX18* 5’UTR region using RT-qPCR (reverse primer in red) after 3D transfection with TBX18 g6 RNA (guide 6) or no transfection (no guide). Regions amplified correspond to #1-5 and are indicated with black arrows. Normalized to beta-actin and relative fold change is compared to no guide.

To test if the depletion of PRC2 results in epigenetic modifications indicative of transcription activation, we performed ChIP-qPCR using H3K27ac and p300 antibodies. Importantly, we observed recruitment of p300 and acetylation mark (H3K27ac) at *TBX18* g6 locus (**Fig 3E**), indicative of active transcription. Another indication of active transcription based on altered PRC2 activity is the recruitment of RNA Polymerase II (Pol II). Therefore, we tested Pol II occupancy at the g6 locus with and without EBdCas9/g6 activity. ChIP-qPCR of Pol II CTD and Pol II CTD Ser5P (indicative of RNA Pol II Pause^49^) revealed a significant recruitment of RNA Pol II to *TBX18* g6 locus due to EBdCas9 activity (**Sup 3A**). RNA Pol II was not only highly enriched in *TBX18* g6 locus at 3D timepoint, but also in 5D timepoint, 2 days after the EBdCas9 was eliminated (**Fig 3F**). Since RNA Pol II is recruited to g6 region, we tested if *TBX18* nascent RNA is transcribed in that region. We tiled no guide and *TBX18*/g6 total RNA with reversed primers targeting *TBX18*5’ UTR region and observed increased nascent RNA amplification in EBdCas9 cells transfected with *TBX18/* g6 compared to no guide using RT-qPCR (**Fig 3G–H).** As controls we validated *TBX18* transcription upregulation on exon 1, but not on 2.0kb upstream region. *TBX18* utilizes two TSS’s at its promoter region (UCSC Genome Browser GRCh37/hg19). In accordance with the tiling experiment, we show EBdCas9/g6 treatment activating transcription in TSS#2 region, 532bp downstream of g6 (**Fig 3G–H**). We show that based on the CAGE-seq data from the FANTOM project, TSS#2 is the dominant promoter/TSS for *TBX18* (**Sup 3B–C**).

### EBdCas9 reveals distal TATA box function

Since *TBX18*/g6 targeted site was highly receptive to transcription activation, epigenetic remodeling and epigenetic memory, we analyzed the chromatin region for 3 possible regulatory DNA sequence elements: G-quadruplex patterns, transcription factor binding site occupancy and core promoter sequence elements. The first tier of these elements is **Q** uadruplex forming **G**-**R** ich **S** equences (QGRS)^50^. G-quadruplex structure motifs have been linked to increased transcriptional activity ^51^. We used the QGRS mapper and identified 10 QGRS repeats on *TBX18* promoter region downstream of g6 locus (**Sup 4A**). The second tier is prediction of transcription factor binding sites, where we identified sparse transcription factors downstream of g6 region and high accumulation of these transcription factors at TSS #1 and TSS#2 (**Sup 4B**). The last tier of these components was identified by the promoter and transcriptional ElemeNT prediction tool ^52^ where potential core promoter elements are detected on *TBX18* promoter region around 1000bp upstream from TSS. These sequences, a combination of TATA box and mammalian initiator factor binding sites (INR) were found downstream of *TBX18* g6 locus (**Fig 4A** and **Sup 4C;** TATA box 21 bp and INR 48-57 bp from g6). Therefore, we challenged the question whether a distal TATA box is functional for *TBX18* transcription by precisely deleting 40bp TATA box (TATATGAC) region, using CRISPR/Cas9-HDR (**Fig 4B**). Two independent, pluripotent EBdCas9 cell lines with homozygous TATA box deletions were generated, TATA Δ #1 and TATA Δ #2 (see materials and methods for details) (**Fig 4B**) and analyzed for *TBX18* transcription. A significant increase of *TBX18* transcription in wild type EBdCas9/g6 (WT) was again confirmed. However, TATA Δ #1, and TATA Δ #2 EBdCas9 lines failed to upregulate *TBX18* transcription (**Fig 4C**). Similar levels of EBdCas9 mRNA expression after Dox induction was observed in all cell lines (**Fig 4C**). From these data we concluded that distal, upstream TATA box is essential for *TBX18* gene expression.

**Figure 4.**
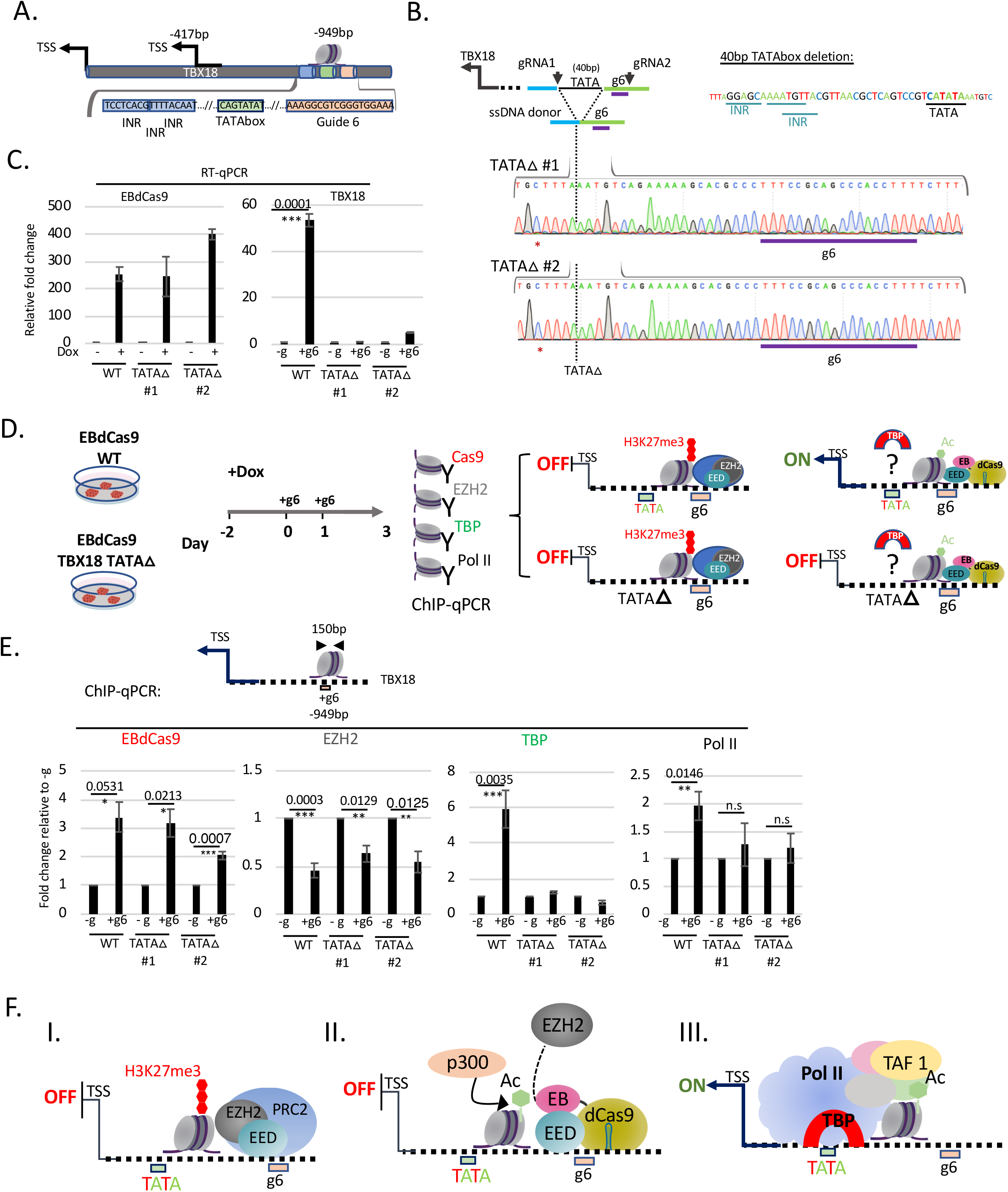
EBdCas9 reveals functional distal TATA box. **A**. TATA box and initiator(INR) representation with respect to guide 6 locus using Element Navigation Tool for detection of core promoter elements. **B**. Generation of TATA box deletion (TATA Δ) clones: TATA Δ #1 and TATA Δ #2 using CRISPR/Cas9. gRNA1 and 2 excise an 88bp fragment, including g6 region. g6 region was reconstructed using 168bp ssDNA donor. Depicted here is a deletion of 40bp including the TATA box and 3 Initiator elements (INR). Genomic DNA of TATA Δ #1 or TATA Δ #2 displaying homozygous clones of 40bp TATA box deletion using Sanger sequencing. PAM site blocking mutation is denoted with a red asterisk. **C**. RT-qPCR of EBdCas9 and TBX18 relative fold change after 3 days Dox induction and TBX18 g6 RNA transfection; normalized to beta-Actin and compared to no guide of each respected cell line (WT- EBdCas9, TATA Δ #1, TATA Δ #2). **D**. EBdCas9 WT and EBdCas9 TATA Δ #1 and TATA Δ #2 timeline for Dox induction, gRNA transfection and ChIP-qPCR analysis using the antibodies Cas9, EZH2, TATA Binding Protein (TBP) and Pol II and analyzing for TBX18 g6 DNA region. Right panel depicts 4 possible model scenarios for TATA box and TATA Δ with and without EBdCas9. **E**. ChIP-qPCR of induced (+Dox) EBdCas9 (WT) and EBdCas9 TATA Δ #1 and TATA Δ #2 3D after transfection with TBX18 g6 RNA (+g6) or no transfection (-g). Normalized to fractioned Input /H3 and compared to −g relative fold change. Antibodies that were used for ChIP are listed above the graphs (Cas9, EZH2, TBP and Pol II) and the genomic region analyzed by qPCR includes *TBX18* g6 locus. \**p*<0.05, ** *p*<0.01, \*\*\**p*<0.001 one-way ANOVA performed. N=2 biological replicates. **F**. Model describing EBdCas9/TBX18 mode of operation: I. *TBX18* distal TATA box function is repressed by PRC2. II. EBdCas9 de-represses *TBX18* TATA box region. III. TBP, TBP Associated Factors (TAF) and Pol II are recruited to *TBX18* TATA box region to allow *TBX18* transcription.

To explore why deletion of distal TATA box affects *TBX18* gene expression, we examined proteins that are recruited to TATA box and allow gene expression. In general, TATA box is a key to core promoter function ^53^. Functional TATA box recruits TATA Binding Protein (TBP), TBP Associated Factors (TAF) and RNA Pol II ^54^ for transcription initiation. If the TATA box, distal from *TBX18* TSS can act as a functional TATA box when PRC2 action is eliminated in this precise location, we expected to identify TBP in the location after EBdCas9 action. To test this, we performed ChIP-qPCR assay and tested whether Cas9, EZH2, TBP and Pol II are located at the g6 region after EBdCas9/g6 treatment (**Fig 4D**). ChIP-qPCR of WT EBdCas9 resulted in EBdCas9 (Cas9) recruitment to g6 site and reduction of EZH2 protein at that site, as seen earlier (**Fig 4E, Fig 2C**). Importantly, ChIP-qPCR of WT EBdCas9/g6 also resulted in recruitment of TBP to g6 region, proving that TATA box 21bp away from g6 region and 900bp away from TSS1 can become a functional TATA box after epigenetic changes (**Fig 4E)**. RNA Pol II was also recruited to g6 region as seen before (**Fig 4E, Fig 3F**, **Sup 3A**). However, when TATA box region was deleted (TATA Δ #1 and TATA Δ #2 EBdCas9), both TBP and Pol II failed to be recruited to TATA box region (**Fig 4E)**. Notably, TATA Δ #1 and TATA Δ #2 EBdCas9 were recruited to g6 region (Cas9) and EZH2 was depleted at guide6 region (**Fig 4E)**. These data show that TATA box >500bp away from *TBX18* TSS acts as a functional TATA box since TBP is recruited to TATA box when PRC2 action is eliminated in the precise locus.

These data suggest that activation of distal TATA box requires the reduction of PRC2 activity in a precise locus. Targeted, localized reduction of EZH2 and H3K27me3 leads to increase of H3K27ac which are recognized by TATA box binding protein associated factor 1 (TAF1) bromodomain to form interactions with TBP and position Pol II for transcription initiation (**Fig 4F**) ^55–60^. These data show that utilizing EBdCas9/g6 identified a PRC2-masked, functional TATA box upstream of the *TBX18* TSS (949pb from TSS#1, 532bb from TSS#2). The TATA box activation requires PRC2 elimination since exclusion of EZH2 in that locus increases acetylation marks that help recruiting TAF1/TBP/Pol II complex and activate *TBX18* transcription.

### EBdCas9 activates *CDKN2A* by epigenetic remodeling

To test for EBdCas9/g effect on PRC2 function in another gene locus, we explored *CDKN2A* (p16) gene epigenetic regulation. P16 is a critical regulator of cell division and a tumor suppressor, that inhibits cyclin D-dependent protein kinase activity and by that reduces G1-S transition^61–64^. In rapidly dividing cells, such as in diffuse intrinsic pontine glioma (DIPG), p16 is repressed due to hypermethylation at the promoter area^63^. We searched for critical loci in p16 gene upstream region to identify potential nucleation sites that would be responsive to PRC2 reduction by EBdCas9. Potential induction of p16 gene by EBdCas9 could suggest new routes for glioma treatment. We therefore tiled the promoter area and gene body of p16 with eight gRNAs ranging from 0.3kb to 2.3kb upstream of TSS and 0.2kb to 0.7kb downstream of TSS (**Fig 5A**). WTC EBdCas9 or NCdCas9 were induced prior to transient transfection of the gRNAs followed by cell harvest at 3D and p16 transcript analysis (**Fig 5B**). EBdCas9 activated p16 transcript expression on 6 out of the 8 gRNAs tested, but none activated transcription in the control, NCdCas9 expressing line (**Fig 5C**). As observed with *TBX18* tiling, gRNAs that are in 0.5kb-1.5kb proximity to TSS showed the highest p16 transcript activation. g1, g2, g3, g4, g6, and g7 induced transcription of p16 ranging from 20-80 fold compared to no guide (-g) or NCdCas9 (**Fig 5A–C**). However, g5 which is 2.2kb upstream of TSS or g8 which is 0.1kb downstream of TSS did not result in significant transcriptional increase, even though EBdCas9/g had access to the DNA in these regions (**Fig 5A–C**). Using Element Navigation Tool^52^ we identified TATA box and mammalian initiator factor binding sites in p16 −424bp and −750bp promoter region, upstream from TSS (**Fig 5D**). These elements are in close proximity to g1, the locus that defines the most powerful PRC2 for regulating p16 transcription. The abundance of regulatory elements predicted in p16 promoter region may be responsible for highly permissive transcript upregulation associated with EBdCas9/gRNA.

**Figure 5.**
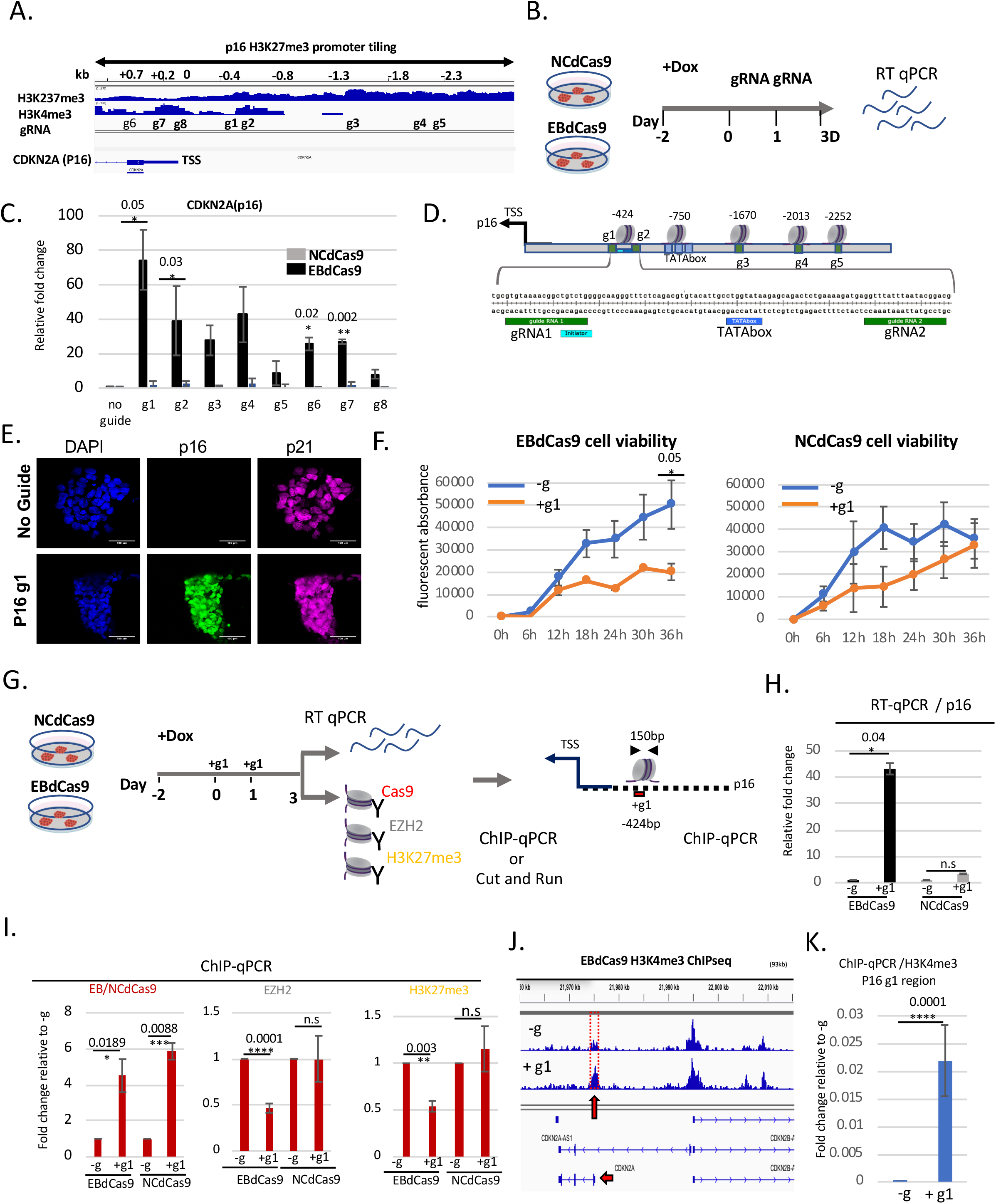
EBdCas9 targeted to specific loci can upregulate CDKN2A (p16) expression. **A**. Integrative Genomic Viewer of *CDKN2A* H3K27me3 and H3K4me3 promoter tiling. **B**. Timeline of EBdCas9 or NCdCas9 induction and gRNA transfection. **C**. RT-qPCR analysis of EBdCas9 or NCdCas9 after 3D transfection with single gRNA (g1-g8) normalized to beta actin and relative fold change compared to no guide (-g). **D**. TATA boxes/initiator representation with respect to surrounding guides loci using Element Navigation Tool for detection of core promoter elements Immunofluorescent imaging of EBdCas9 after 3D transfection with single p16 gRNA 1 (+ g1) or no transfection (-g). Blue-Dapi, Green-p16, Far red-p21; scale bar is 100μm. **F**. Cell viability of EBdCas9 and NCdCas9 transfected with p16 g1 (+g1) compared to no guide (-g). Times points are every 6h measured by fluorecin using Alamar Blue. **G**. EBdCas9 timeline for measuring epigenetic remodeling. **H**. RT-qPCR of p16 relative fold change of EBdCas9 and NCdCas9 after 3D dox induction and p16 g1 transfection. Samples were normalized to beta-Actin and compared to no guide (-g) of each respected cell line. **I.** ChIP-qPCR of induced (+Dox) EBdCas9 and NCdCas9 after 3D transfection with p16 g1 RNA (+g1) or no transfection (-g). Normalized to fractioned Input/ H3 and compared to −g relative fold change. Antibodies that were used for ChIP are listed above the graphs (Cas9, EZH2, H3K27me3) and the genomic region analyzed by qPCR includes p16 g1 locus. \**p*<0.05, ** *p*<0.01, \*\*\**p*<0.001 one-way ANOVA performed. n=3 biological replicates. **J-K.** H3K4me3 marks increase after EBdCas9/p16 g1 (+g1) induction. Cut and Run analysis (**J**) or ChIP-qPCR (**K**) of induced (+Dox) EBdCas9 after 3D transfection with p16 g1 RNA (+g1) or no transfection (-g) for H3K4me3. H3K4me3 tracks are normalized to IgG and displayed on Integrated Genome Viewer (IGV); 93kb CDKN2A (p16) window viewer. Red dots demonstrate CDKN2A 2.5kb of TSS region. ChIP-qPCR region primer sets cover region g1 as displayed in G. normalized to fractioned Input/ H3 relative to no guide (-g). n=3 biological replicates.

We showed that p16 was also upregulated on protein level by using immunofluorescence analysis (**Fig 5E**). Since activation of p16 expression results in cell cycle arrest in gliomas^63^, we tested the effect of p16 upregulation in iPSC. Transfection of WTC EBdCas9 with g1(p16) resulted in 2 fold cell and colony size reduction compared to no guide controls (-g) 3 days post transfection (**Sup 5A–B**). In agreement, EBdCas9/g1(p16) resulted in poor cell viability compared to no guide (-g) or NCdCas9 using resazurin fluorescent (Alamar Blue) detection reagent (**Fig 5F)**.

Tracing of EBdCas9 and NCdCas9 proteins in g1 locus of p16 gene by ChIP-qPCR showed equivalent protein loading to the chromatin locus, however, EBdCas9/g1, but not NCdCas9/g1 resulted in 40 fold transcriptional upregulation of p16 gene (**Fig 5G–H**). Similarly, EBdCas9/g1, but not NCdCas9/g1 resulted in significant reduction of EZH2 protein and H3K27me3 marks at targeted chromatin site, analyzed using ChIP-qPCR (**Fig 5I**). Accumulation of H3K4me3 marks at TSS of EBdCas9/g1 compared to -g samples is indicative of active transcription, analyzed using CUT and RUN method^65^ and validated by ChIP-qPCR (**Fig 5J–K**). ChIP-qPCR of a distant region (−2300bp form TSS) that does not lead to gene activation does not show reduced EZH2 levels or H3K27me3 marks either, suggesting possible steric hindrance between EBdCas9 and PRC2 accessibility (**Sup 5C–D**).

Similar to that seen with *TBX18*, these data suggest that targeted EBdCas9/g1 identified a masked TATA box ~400bp upstream of the TSS of p16 gene. We propose that in the case of p16 the upstream TATA box inactivation requires PRC2 activity since eliminating EZH2 in the proximity of the TATA box promotes p16 transcription.

### EB expression induces trophoblast fate

The first lineage bifurcation, trophoblast vs ICM cellular fate decision is dependent on PRC2^36^. While overexpression of H3K27me3 is associated with ICM lineage, genome wide depletion of H3K27me3 marks is associated with trophectoderm lineage^3, 36, 66–69^(**Fig 6A**). We therefore tested if the EED binder (EB; ^35^) is able to induce trophoblast fate (**Fig 6A)**. Recently two groups have generated culture condition that enabled the establishment of extended pluripotent stem cells (EPS) from either cleavage state of mouse embryos or human embryonic stem cells^67, 68^. The EPS cell stage has a developmental potency and capability to the first cell fate bifurcation to generate both embryonic, inner cell mass (ICM) and extraembryonic placental tissue, as trophectoderm (TE) cell lineage^67, 68^. Moreover, EPS epigenetic analysis validated enrichment of bivalent, H3K27me3 and H3K4me3 marks in developmental ^67, 68^. We therefore reprogrammed WTC EB-Flag and WTC NC-Flag^35^ to EPS stage using LCDM (hLIF, chir99021, (S)-(+)-dimethindene maleate:DiM, and minocycline hydrochloride: MiH) reprogramming cocktail 68. Once established, we validated colony dome-shaped morphology, single cell colony efficiency and expression of pluripotency markers (**Fig 6B)**. Validated EPS(EB-Flag) and EPS(NC-Flag) were assayed for TE differentiation to determine whether reduction of H3K27me3 marks using EB accelerated TE lineage choice (**Sup 6A**). EPS(EB-Flag) and EPS(NC-Flag) were grown on matrigel (+Dox) in trophoblast differentiation media (TX) containing TGFb, FGF4 and heparin^70^. EB-Flag, but not NC-Flag expressing cells lost EPS colony morphology at 4D, suggesting accelerated differentiation (**Fig 6C)**. Not only *GATA3* and *TBX3* mRNA expression validated accelerated differentiation compared to uninduced cells but *OCT4* was also significantly reduced (**Fig 6D**).

**Figure 6.**
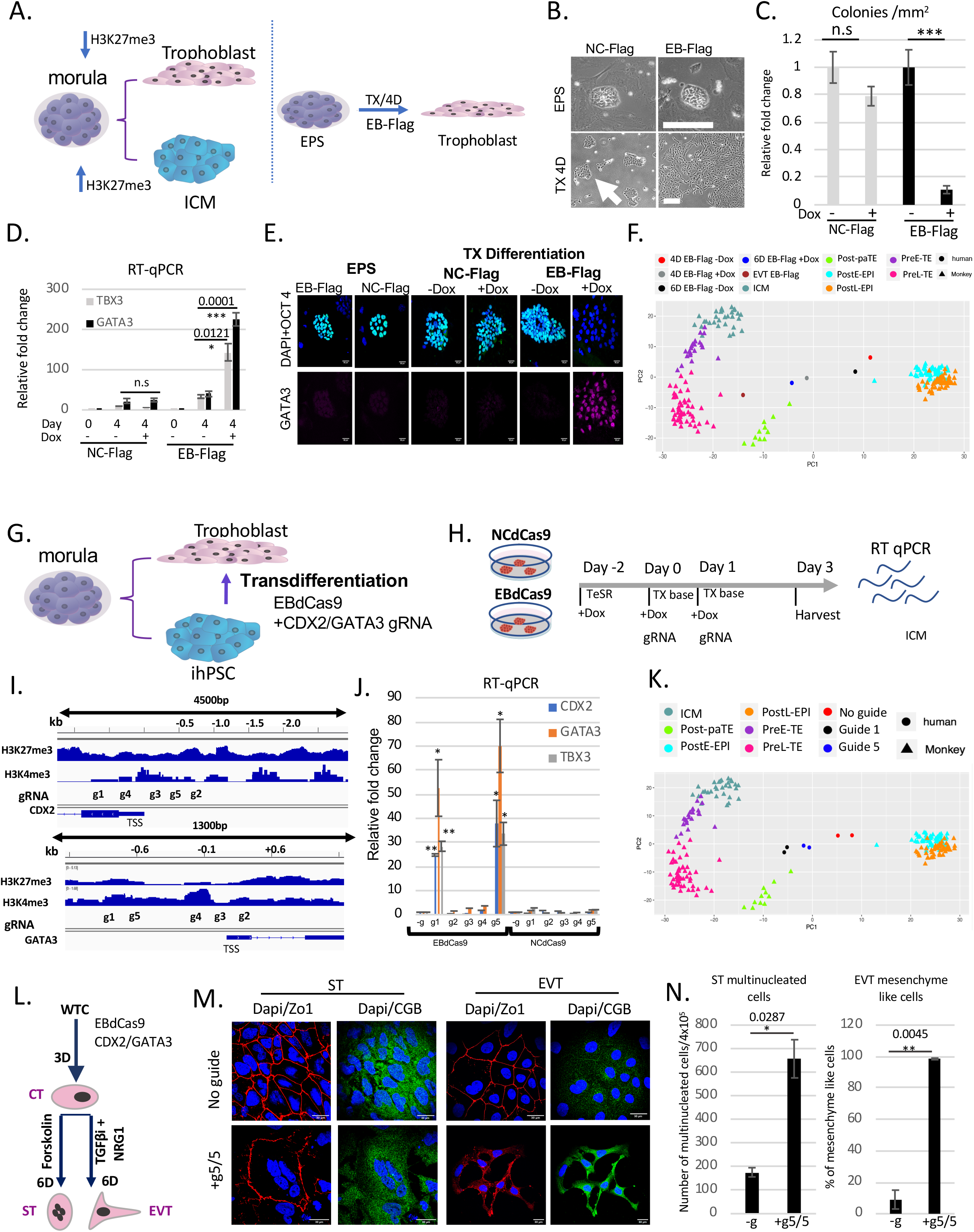
Trophoblast trans-differentiation using EBdCas9. **A**. Morula bifurcation to trophoblast and ICM is H3K27me3 dependent (left panel). EPS differentiation using EB-Flag and TX media ^70^ (right panel). **B**. WTC EB-Flag or WTC NC-Flag as described previously ^35^ were reprogramed to EPS for 2 weeks (3 passages). Bright Field of colony morphology of EPS EB-Flag or NC-Flag induced with Dox on MEF for 4D or plated on matrigel in TX media (TGFb, FGF4 and heparin) for 4 days (TX 4D). **C**. Colony morphology count following 4D in TX media EB-Flag compared to NC-Flag. **D**. RT-qPCR of EB-Flag and NC-Flag either in EPS stage or 4D in trophoblast differentiation (TX media) with (+) or without (−) Dox. Normalized to beta-actin and relative to EPS EB-Flag stage**. E**. Immunofluorescence of EPS EB-Flag or EPS NC-Flag on MEF / LCDM media or differentiation using matrigel/ TX media with (+) or without Dox (−). Dapi-Blue, WGA-Red, Oct4- Green, Gata3- Far red. Scale bar is 50μm. **F**. PCA analysis of EPS samples compared to monkey single cell RNA-seq ^73^. EB-Flag EPS cells were differentiating in TX media with or without dox for 4 days or 6 days or passage 3 times as extravillous cytotrophoblast (EVT). Cell types in the monkey single cell data include: Post-paTE, post-implantation parietal trophectoderm; PreL-TE, pre-implantation late TE; PreE-TE, pre-implantation early TE; ICM, inner cell mass; Pre-EPI, pre-implantation epiblast; PostE-EPI, post-implantation early epiblast; PostL-EPI, post-implantation late epiblast. **G**. Model of WTC EBdCas9 trans-differentiation to trophoblasts using CDX2 and GATA3 gRNA cocktail. **H**. Timeline of EBdCas9 or NCdCas9 induction and gRNA transfection. **I**. Tiling of CDX3 and GATA3 promoter and gene body gRNA. **J**. RT-qPCR analysis of CDX2 and GATA3 of Co-transfected gRNAs in the presence of EBdCas9 or NCdCas9 induction (+Dox). g1-g5 corresponds to g1/g1, g2/g2, g3/g3, g4/g4, g5/g5 CDX2/GATA3 co-transfection. Normalized to beta-actin and compared to no guide (-g). **K**. PCA analysis of 3D WTC EBdCas9 co-transfected with g5/g5 CDX2/GATA3 and g1/g1 CDX2/GATA3 or not transfected (-g) bulk RNAseq compared to monkey single cell RNA-seq ^73^. Cell types are exactly as in (F). **L**. Timeline of ExtraVillous Trophoblasts(EVT) and SyncytioTrophoblast(ST) differentiation after EBdCas9:CDX2/GATA3 g5,g5 transfection(cytotrophoblast (CT) like stage) and the factors used. **M**. Immunofluorescence of EBdCas9 3D post CDX2/GATA3 g5,g5 RNA transfection compared to no guide, and further 6 days (6D) differentiation to either EVT using 7.5mM TGFbi and 100ng/ml (NRG1) or ST using 2mM forskolin. Dapi- blue,ZO-1-red and CGB (chorionic gonadotropin beta) −green. Scale bar is 30μm. **N**. Quantification of ST (multinucleation) and EVT (mesenchyme like morphology) positive cell count. Area of total count is 18mm^2^. N=2 for biological replicas.

Confocal imaging confirmed the tight, dome-shaped morphology, expression of nuclear stem cell transcription factor OCT4 and absence of GATA3 expression for both EPS(EB-Flag) and EPS(NC-Flag) at day 0 (**Fig 6E**). However, TE differentiation (4D) with EB-Flag, but not NC-Flag resulted in dramatic reduction of OCT 4 expression, loss of colony morphology and upregulation of GATA3 (**Fig 6E**). Furthermore, immunoblot analysis revealed that 4D TE differentiation with EB-Flag reduced H3K27me3, EZH2 and Oct4 while GATA3 was upregulated (**Sup 6B)**.

To investigate these fate changes in more detail we analyzed gene expression in these samples utilizing RNA seq, as done before ^71^. We projected TE differentiated EPS(EB-Flag) cells (4D, 6D and differentiated ExtraVillus Cytotrophoblast (EVT) ^72^) onto the principal component analysis (PCA) plot of single cell transcriptomes of early cynomolgus monkeys^73^ and found acceleration of TE differentiation in EB-Flag expressing cells (**Fig 6F**). The projection is based on 773 highly variable genes (standard deviation>2) in the monkey single cell dataset. TE differentiated EPS EB-Flag cells that were induced with Dox during differentiation to express EB-Flag emigrate from post-implantation early or late epiblast (PostE-EPI ; PostL-EPI), to post-implantation partial trophectoderm (Post-paTE) and pre-implantation late trophectoderm (PreL-TE) compared to no Dox EB-Flag differentiated cells. Also, the PCA clearly showed that all EB-Flag samples are distant from ICM on the TE differentiation lineage, showing that EB-Flag accelerates TE differentiation. EVT EB-Flag cells were first induced to express EB for 4 days and thereafter passaged 3 times (in TSC conditioned media) ^72^ without Dox (and therefore without further EB expression) and found to be closest to pre late trophectoderm (PreL-TE). Overall, these results confirm that elimination of H3K27me3 marks by induction of the EB-Flag protein dramatically accelerates TE lineage differentiation.

### Targeted PRC2 inhibition on *CDX2* and *GATA3* using EBdCas9 induce trophoblast trans-differentiation

During mouse blastocyst formation, the relative levels of EED and KDM6B determine altered PRC2 complex recruitment and incorporation of H3K27me3 marks at the chromatin domains of target genes (**Fig 6A**). Two trophectoderm (TE) lineage-specific transcription factors CDX2 and GATA3 show PRC2 dependent repression in the ICM^36, 37^. However, it is not known if PRC2 activity is sufficient in the promoter regions of these two genes to distinguish the bifurcation between TE and ICM lineages. We tested if ICM like cells, such as iPSC, which have already passed the bifurcation point, are able to transdifferentiate to TE if CDX2 and GATA3 H3K27me3 epigenetic marks are reduced (**Fig 6G**). To do so, we targeted EBdCas9/gRNA to the gene-upstream regions of TE transcription factors *CDX2* and *GATA3*. WTC EBdCas9 was grown on matrigel in TeSR (+Dox) for 2 days, and after gRNA transfection, the media was changed to TX base media (+Dox) without factors (no TGFb, FGF4 and heparin) (**Fig 6H**). This created a permissive environment for TE differentiation, in which EBdCas9/gRNAs are the sole drivers for potential trans-differentiation. *CDX2* and *GATA3* were tiled across the promoter and gene body region with 5 different guides (**Fig 6I**). Since these two transcription factors are critical players in TE differentiation in mouse^36^, we screened for the critical regions by co-transfected g1 from *CDX2* and g1 from *GATA3* and applied this two-gene co-transfection scheme to all CDX2/GATA3 gRNA combinations (g1/g1, g2/g2,…g5/g5). WTC EBdCas9/gRNA, but not NCdCas9/gRNA cocktail 1 and 5 resulted in a significant gene activation (20-80 fold increase) for *CDX2* and *GATA3*, as well as a TE marker *TBX3* (**Fig 6J, Sup 6C**). gRNA cocktails g(2-4) didn’t show any gene activation to either *CDX2* or *GATA3*.

To investigate WTC-TE trans-differentiation changes in more detail we analyzed global transcriptomics of g1/g1 and g5/g5 CDX2 and GATA3 samples by RNA seq. Projection of WTC EBdCas9 in TX (base) media with or without *CDX2* and *GATA3* g1/g1 or g5/g5 cocktail on PCA of single cell transcriptome of early cynomolgus monkeys ^73^ showed advanced emigration of g1/g1 or g5/g5 from postE-EPI/ postL-EPI towards Post-paTE and PreL-TE compared to no guide control (**Fig 6K**). We also compared the transcriptomics of *CDX2* and *GATA3* cocktail to trophoblast dataset as previously shown ^74^. Similarly, PCA projection of developmental genes showed that WTC EBdCas9 with no gRNA transfection either 3D TeSR or TX (base) conditions associated with control (WTC) dataset. More importantly, transcriptomics of WTC EBdCas9 (TX) transfected with either g1/g1 or g5/g5 cocktail for *CDX2* and *GATA3* corresponded to cell that had differentiated towards TE fate (**Sup 6D**). To further test if WTC EBdCas9 g5,g5 *CDX2* and *GATA3* cocktail is able to produce cytotrophoblast progenitor cells following 3D of trans-differentiation, we proceeded to specific extravillous trophoblast (EVT) or Syncytiotrophoblast (ST) 6 days (6D) differentiation using TGFbi and Neuregulin or Forskolin, respectively ^72^ (**Fig 6L**). Immunofluorescence staining confirmed that 3D WTC EBdCas9 g5,g5 *CDX2/GATA3* cocktail are able to differentiate to EVT and ST due to both positive staining of chorionic gonadotropin beta (CGB) and mesenchyme like and multinucleation morphology respectively^72^ (**Fig 6M–N, Sup 6K**). Overall, we can conclude that EBdCas9 targeting only *CDX2* and *GATA3* epigenetic state is sufficient to trans-differentiate iPSC to trophoblast-like state. These data confirm that targeting EBdCas9 to eliminate H3K27me3 in precise *CDX2* and *GATA3* loci results in trans-differentiation of iPSC to TE.

We also used the EPS to directly test the bifurcation process. Since g1,g1 and g5,g5 RNA resulted in significant *CDX2* and *GATA3* gene activation in WTC EBdCas9 cell lines, we also reprogramed WTC EBdCas9 to EPS state and measure *CDX2* and *GATA3* gene activation in EPS(EBdCas9)/g1,g1 and g5,g5 samples (**Sup 6E–G**). EPS(EBdCas9) based CDX2 and GATA3 gene activation increased between 100-500 fold, reminiscent of EB-Flag TE differentiation gene activation results (**Fig 6D**). ChIP-qPCR analysis of WTC EBdCas9 TE differentiation using g5,g5 gRNA for *CDX2* and *GATA3* cocktail, resulted in EBdCas9 recruitment and reduction of H3K27me3 and EZH2 at targeted genomic locus, compared to -g (**Sup 6H–I**). Additionally, Cut and Run analysis showed reduction of H3K27me3 marks in EBdCas9/g1,g1 or g5,g5 compared to no guide on GATA3 TSS and gene body **(Sup 6J)**. These data suggest that PRC2 activity in precise chromatin locations of only two genes can direct the first bifurcation to ICM fate.

## DISCUSSION

Control of epigenetic regulation holds immense therapeutic promise for human disease without chemical drugs or manipulations of endogenous genes. Here we develop a system for targeted inhibition of PRC2 that identify the precise nucleosomes that require H3K27me3 marks for transcription repression. We identified a distal TATA box that is regulated by PRC2 repression and required for transcript activation. We show that this PRC2 inhibiting tool, EBdCas9, allows the study of epigenetic memory of PRC2 at specific loci. Furthermore, we show the general applicability of EBdCas9, by identifying the regions where PRC2 disruption induces transcription of bivalent genes (*TBX18*, *p16*, *CDX2* and *GATA3*). In total, we have targeted 26 specific sites in promoter regions upstream of four different genes, and observed significant PRC2-dependent transcriptional derepression in 8 loci. NCdCas9, which differs from EB by two amino acids required for EED binding, does not result in transcriptional changes. This suggests that EBdCas9 functions by locally inhibiting PRC2 activity. When targeting PRC2 in the promoter regions of known trophectoderm transcription factors *CDX2* and *GATA3*, EBdCas9 application identified specific loci in which PRC2 elimination is sufficient to upregulate the gene transcription and therefore induce trophectoderm cell fate commitment (**Fig 6**).

Dead-Cas9 (dCas9) is the most promising method for targeted delivery of agents to chromatin. However, previous fusions of dCas9 to enzymes result in off target activity outside of the targeted locus. Existing histone modifying enzyme fusions are continuously active, potentially introducing or removing marks at off target locations independent of the endogenous epigenetic modification machinery. In contrast, the computational designed protein EB fused to dCas9 (EBdCas9) with appropriate gRNA presented here is able to precisely inhibit PRC2 at specific genetic loci, enabling: (1) target and inhibition of PRC2 at a single nucleosome level, (2) reduced H3K27me3 at precise, targeted loci, (3) induced targeted transcription, (4) mediation of neighborhood spreading of remodeled epigenetic marks, (5) generation of epigenetic memory, (6) discovery of functional distal TATA box and TBP >500bp of TSS, (7) change of cell functionality, and (8) transdifferentiation of one cell fate to another. Recently, several groups have activated targeted genes using synergistic activation mediator (SAM) such as dCas9-VP64, sgRNA, and MS2/p65/HSF1^75–78^. Their strategy was to directly overexpress targeted genes of interest by recruitment of transcription factors to the promoter region. Our approach of targeted PRC2 inhibition utilizing EBdCas9/gRNA allows organic expression of targeted genes as the cell makes holistic decisions for transcriptional activation. EBdCas9 function is specific to PRC2 repressed genes.

One of the fundamental findings of this study is that broad H3K27me3 marks on promoter regions appear indiscriminate. Using EBdCas9/gRNA we identified key PRC2 dependent single nucleosomes on promoter regions that control gene expression. In eukaryotes, genes are classified as TATA-containing and TATA-less in their promoter regions ^79^. The TATA-containing core promoter, in a range of −150 to −1 relative to TSS constitute ~17% of the total promoters in yeast, ~10% in worms, ~14% in fruit flies, ~10% in zebrafish and ~3% in human and mouse making 97% of core promoter TATA-less ^80^. If the −500 to +500 region spanning TSS is considered, the number of predicted TATA-containing promoter sequences grows to 14% ^81^. While TATA-containing promoters have TATA sequence at region −30 to −1 relative to TSS region, TATA-less promoters have higher GC content in the TSS −150 to −1 region ^80, 82^. In addition, human TATA-less promoters have different structural motifs such as G-quadruplex (GNGNGNG) and CpG islands that are enriched at approximately ~42% at TATA less promoter regions ^80^. In the scenario of a distal TATA box, we hypothesize that G-quadruplex structure contributes to active transcription as nucleosome-depleted regions in euchromatin associate with H3K4me3, RNA Pol II, and transcription factor recruitment ^51^. Here, we identify single nucleosomes on key promoter regions, that are transcriptionally responsive to EBdCas9, suggesting that these regions are normally repressed due to PRC2 activity. In the case of *TBX18* we show that PRC2 dependent sites co-localize with distal TATA box and initiator binding elements. Furthermore, we show that the TATA box region is required for EBdCas9 based gene upregulation since TBP is recruited to TATA box site after PRC2 de-repression by EBdCas9. CRISPR/Cas9-based deletion of the TATA box region eliminates TBP recruitment which subsequently, results in transcript inactivation. This suggests that remote TATA-boxes are regulated by biological signals culminating in PRC2 activity. We now show that the EBdCas9 biological tool can reveal masked regulatory elements on promoter regions through EBdCas9/gRNA multiplex promoter screens.

The combination of controlled epigenetic gain- and loss-of-function manipulations are the most desirable for elastic gene expression based epigenetic memory. In the future, *in vivo* EBdCas9 expression under tissue specific promoters will allow us to explore the *in vivo* applicability of this technology. The adaptive and efficient targeted PRC2 inhibition by EBdCas9 identifies functional H3K27me3 marks and mediates gene activation, which holds promise both as an epigenetic tool in biomedical research and as an approach for treating a wide range of human diseases.

## Acknowledgements

We thank members of the Ruohola-Baker laboratory for helpful discussions, inspiration and advice throughout this work. We thank Christopher Cavanaugh, Jennifer Hesson Jay Sarthy, Daniel Mar, Infencia Xavier Raj, Pushpa Pushpa and James Moody for help and advice throughout this work. SL was supported by WRF Postdoctoral Fellowship and ISCRM Fellows program. This work is supported in part by grants from the National Institutes of Health R01GM097372, R01GM97372-03S1, R01GM083867, 1P01GM081619 for HRB.

## Author Contributions

S.L. and H.R-B. conceived and designed the experiments. L.S, D.I-C, and H.H performed the experiments and/or analyzed the resulting data. Y.W. and A.A. performed the bioinformatic analysis of the RNA-seq. D.B,R.D.H and K.B provided materials, and advice. S.L and H.R.B. wrote the manuscript with input from the other authors.

## EXPERIMENTAL PROCEDURES

### hiPSC and hESC Cell culture

The hiPSC line **WTC** #11, previously derived in the Conklin laboratory ^83^, was cultured on Matrigel (1:30) growth factor-reduced basement membrane matrix (Corning) in mTeSR media (StemCell Technologies). Cells were passaged using versene once reaching 70% confluency and plated at 1:6 density. For **EPS** conditions, cells were grown as previously described^68^. Briefly, WTC cells were reprogrammed in base medium containing 100 mL DMEM/F12, 100 mL Neurobasal, 1 mL N2 supplement, 2 mL B27 supplement, 1% GlutaMAX, 1% NEAA, 0.1 mM β-mercaptoethanol, penicillin-streptomycin and 5% KSR, and freshly supplemented with 10 ng/ml hLIF, GSK3i (1 μM), ROCKi (2 μM), (S)-(+)-Dimethindene maleate (2 μM; Tocris), Minocycline hydrochloride (2 μM; Santa Cruz Biotechnology) and IWR-endo-1 (0.5-1 μM; Selleckchem). Cells were adapted to EPS conditions for at least 3 passages before analysis. EPS cells were pushed toward differentiation using TX media ^70^: TX medium formulation was DMEM/F12 without HEPES and L-glutamine (Life Technologies), 64 mg/l l-ascorbic acid-2-phosphate magnesium, 14 mg/l sodium selenite, 19.4 mg/l insulin, 543 mg/l NaHCO3, 10.7 mg/l holo-transferrin (all Sigma-Aldrich), 25 ng/ml human recombinant FGF4 (Reliatech), 2 ng/ml human recombinant TGF-ß1 (PeproTech), 1 μg/ml heparin (Sigma-Aldrich), 2 mM L-glutamine, 1% penicillin, and streptomycin (all PAN-biotech). Medium was prepared without growth factors (TX-growth factors) and stored at 4° C. To prepare complete TX, the growth factors: FGF4, heparin, and TGF-ß1 were added prior to use. Medium was changed every other day. All cells were cultured at 37° C in 5% CO_2_. For **Elf1 iCas9** conditions, cells were treated as described before^46^ but adapted to TeSR culture media and single cell passaging: the cells were grown on Matrigel (1:30) growth factor-reduced basement membrane matrix (Corning) in mTeSR media (StemCell Technologies) and passaged using versene once reaching 70% confluency and plated at 1:6 density.

### EBdCas9 and EBNCdCas9 plasmid construction

we used the AAVS1 TREG KRAB-dCas9 plasmid previously derived in the Conklin laboratory ^83^, eliminated KRAB using PacI and AgeI restriction cut and then constructed and religated the EEDbinder-linker-dCas9-NLS-mCherry (EBdCas9) or EEDbinder Negative Control-linker-dCas9-NLS-mCherry (EBNCdCas9) to the cut plasmid, screened colonies and verified the sequence by Sanger sequencing (see supplemental material for exact sequence).

### Insertion of inducible EBdCas9 and EBNCdCas9 into AAVS1 site of WTC cells

1×10^6^ cells of WTC p42 were transfected with 5mg AAVS1-TALEN R plasmid (Addgene #59026), 5 μg AAVS1-TALEN L plasmid (Addgene #59025), and 5mg donor plasmid (AAVS1 TREG EBdCas9 or AAVS1 TREG EBNCdCas9) using the Amaxa Lonza Human stem cell Kit #2. The cells were then plated with 5 μM of Rock inhibitor (ROCKi) onto 10cm with fresh media. Three days following the nucleofection, the cells were selected for neomycin resistance with Genetecin (50mg/ml) for four days. Clones that survived after selection were expanded as a pool. The clones were plated onto matrigel with or without doxycycline (2 μg/ml) and RNA was extracted in order to analyze the level of Cas9 expression by qPCR. Insertion of EBdCas9 or NCdCas9 into the AAVS1 site was confirmed by cellular genomic isolation, PCR amplification and Sanger sequencing.

### Trophoblast differentiation using EB-Flag/NC-Flag EPS cells

50k-100k EPS Cells grown on matrigel (1:30) coated plate in TX base medium (DMEM/F12 w/o HEPES or L-Glu (21331-020), 64mg/ml ascorbic acid phosphate Mg, 14ug/l sodium selenite, 19.4mg/l insulin, 10.7 mg/l holo-transferrin, 543mg/l NaHCO3, 1% Pen Strep and, 2mM Glutamax) and TX factors (25ng/ml FGF4, 2ng/ml TGF-beta1and 1 mg /mL Heparin) TX media was changed every other day. 2mg/ml doxycycline was added to the medium on day of differentiation.

### Trophoblast trans-differentiation using EBdCas9/NCdCas9 WTC cells

100k WTC cells were plated on matrigel plates (1:30) in the presence TeSR and doxycycline (2mg/ml) for 48h. Prior to gRNA transfections the media was changed to TX media as described in derivation and maintenance of murine trophoblast stem cells under defined conditions^70^ (without factors). gRNAs were transfected using RNAimax for 2 consecutive days and were either harvest for RT-qPCR, ChIP-qPCR at day 3 or continued for further EVT/ST differentiation using the below media conditions for EVT or ST, without EB or NC induction.

### EVT/ST differentiation

3d EBdCas9 TX cells were grown to 80% confluency in TX medium ^70^ and dissociated with 0.05% Trypsin-EDTA for 5 min at 37° and harvested as described in derivation of human trophoblast stem cells^72^. Briefly, trypsinization was stopped using EVT/ST base medium (DMEM/F12 0.1 mM 2 mercaptoethanol, 0.5% Penicillin-Streptomycin, 0.3% BSA, 1% ITS-X supplement, 2.5 mM Y27632 (Rock i), 4% KnockOut Serum Replacement) and cells were plated on matrigel (1:30) 0.75×10^5^ density per well and cultured in 2 mL of EVT/ST medium with either EVT differentiation factors (100 ng/ml NRG1, 7.5 mM TGF-ßi A83-01) or ST differentiation factor (2mM forskolin). Cell were cultured for additional 6 days, with media changes every other day. On day 9 cells were fixed and permeabilized for immunofluorescent staining and imaging.

### Guide RNA design, synthesis and transfection

The gRNAs targeting TBX18, P16, CDX2 and GATA3 genes were designed using the CRISPRscan web prediction tools ^45^ and ordered as T7-gRNA primers. The T7-gRNA forward primer and the reverse scaffold primer were used in primer extension reaction to synthesized a double stranded DNA fragment by using Q5 High Fidelity-based PCR (New England Biolabs) followed by PCR purification (Qiagen). The 120 bp dsDNA served as a template for IVT (MAXIscript T7 kit, applied Biosystems). The RNA was then purified using Pellet Paint® Co-Precipitant (Novagen). For transfection, WTC EBdCas9 or NCdCas9 cells were seeded at day 0, and treated with doxycycline (2μg/ml) for 2 days before and during transfection. On day 2 cells were transfected with gRNAs using Lipofectamine RNAiMAX (Life Technologies). gRNA was added at a 40□nM final concentration when added alone or 20nM in co-gRNA transfection. A second transfection was performed after 24□h. Two days after the last gRNA transfection, cells were harvest for either DNA, RNA, protein, ChIPqPCR, or Cut and Run analysis.

### Elf iCas9 transfection

Elf iCas9 cell line^46^ was induced with dox for 2 days prior to gRNA transient transfection. individual gRNAs were transfected transiently (800ng/ml) and cell were collected for genomic DNA isolation, amplification of designated regions and indel validation using Sanger sequencing.

### DNA extraction and sequencing

Genomic DNA was collected using DNAzol reagent (Invitrogen) according to manufacturer’s instructions and quantified using Nanodrop ND-1000. Genomic regions flanking the AAVS1 were PCR amplified with the designed primers, purified by PCR Purification Kit (Invitrogen) and sent to Genewiz for sequencing.

### RNA extraction and RT-qPCR analysis

RNA was extracted using Trizol (Life Technologies) according to manufacturer’s instructions. RNA samples were treated with Turbo DNase (ThermoFischer) and quantified using Nanodrop ND-1000. Reverse transcription was performed using iScript (BioRad). 10 ng of cDNA was used to perform qRT-PCR using SYBR Green, with suitable primers on an Applied Biosystems 7300 real time PCR system with PCR conditions as stage 1 50°C for 2mins, stage 2 as 95°C for 10mis, 95°C for 15sec, 60°C for 1min(40 Cycles). ß-actin was used as an endogenous control (Sup Table 4).

### Protein extraction and Western blot analysis

Cells were lysed directly on the plate with lysis buffer containing 20mM Tris-HCl pH 7.5, 150mM NaCl, 15% Glycerol, 1% Triton x-100, 1M *ß*-Glycerolphosphate, 0.5M NaF, 0.1M Sodium Pyrophosphate, Orthovanadate, PMSF and 2.3% SDS. 25 U of Benzonase® Nuclease (EMD Chemicals, Gibbstown, NJ) was added to the lysis buffer right before use. Proteins were quantified by Bradford assay (Bio-rad), using BSA (Bovine Serum Albumin) as Standard using the EnWallac Vision. The protein samples were combined with the 4x Laemli sample buffer (900 μl of sample buffer and 100 μl β-Mercaptoethanol), heated (95°C, 5mins) and run on SDS-PAGE (protean TGX pre-casted gradient gel, 4%-20%, Bio-rad) and transferred to the Nitro-Cellulose membrane (Bio-Rad) by semi-dry transfer (Bio-Rad). Membrane was blocked for 1hr with 5% milk, and incubated in the primary antibodies overnight in 4°C. The antibodies used for western blot were beta-Actin (Sigma A5441, 1:2000), Cas9 (Cell Signaling 1:1000), Oct-4 (Santa Cruz sc-5279, 1:1000, Novus Biologicals NB110-90606, 1:500), H3K27me3 (Active Motive 39155 1:1000), EZH2 (Cell Signaling D2C9, 1:1000), GATA3 (Cell signaling D13C9,1:1000), Flag (sigma F1804, 1:1000)The membranes were then incubated with secondary antibodies (1:10000, goat anti-rabbit or goat anti-mouse IgG HRP conjugate (Bio-Rad) for 1hr and the detection was performed using the immobilon-luminol reagent assay (EMP Millipore).

### Immunostaining and confocal imaging

Cells were fixed in 4% paraformaldehyde for 15 min, washed with PBS (3×5min), and blocked in the presence of 0.1% Triton X-100 and 2% BSA for 1h at room temp. Cells were then incubated in primary antibody overnight, washed with PBS (3×5min), incubated with the secondary antibody in 2% BSA for 1hr, washed (4×10mins, adding 1mg/ml DAPI at the 2nd wash), mounted using Vectashield^®^ (Vector Labs) and imaged using a Leica TCS-SPE confocal microscope at 10x, 40x or 63x. Antibodies use in immunostaining were: TBX18 (Santa Cruz sc-514486, 1:200), Oct 4 (Novus Biologicals, NB11- 1:150), P16 (Invitrogen MA5-14260, 1:250), p21 (Cell Signalling 2947S, 1:200), GATA3 (Cell signaling D13C9, 1:250), CGB (DAKO GA508; 1:200). Slides were rinsed with PBS-T and incubated with secondary antibodies: DAPI (0.02 μg/mL, Molecular Probes), goat anti-rabbit 488, and goat anti-mouse 647 (1:500, Molecular Probes) for 2 hours at room temperature in blocking buffer. (EBdCas9 or NCdCas9 were visualized using endogenous mCherry expression).

### Cell harvesting Cross-linking and PIXUL sonication

All PIXUL steps began from 1×35 mm plate. Cells were harvested using Versene (Thermo Fisher), followed by 3 min centrifugation (1.2 xg). Pelleted cells were washed with 1X PBS. Cells were cross-linked by adding 500ul 1% formaldehyde in PBS for 20min in RT. Supernatant was removed and 500ul of PBS/glycine (125mM) was added for 5 min at RT for quenching. Supernatant was removed and the cells were washed with 500ul of PBS. PBS was removed and samples were stored in −80°C. For shearing, cells were resuspended in 100ul chromatin sheering buffer (Active Motif) and placed on 96 well plate sealed with PCR film (MiniAMP Optical Adhesive Film) and placed in PIXUL for shearing. PIXUL was programmed to sonicate for 50 pulse at 1.00 (kHz) PRF (pulse repetitive frequency) for 4.0 min at 20.00 burst rate (Hz). 6 samples were sheered at a time for 4 times at the same settings. Once sheered the cells were chromatin immunoprecipitated using matrix-ChIP.

### Matrix-ChIP-qPCR analysis

Matrix-ChIP was performed as previously described^47, 84^. Briefly, UV treated 96-well polypropylene microplates were incubated with protein A on a low-speed shaker at room temperature overnight. The next day, the wells of the 96-well plate-coated with protein A were blocked with blocking buffer containing 5% BSA, sheared salmon sperm DNA (10 μg/ μl final) in immunoprecipitation buffer (150 mM NaCl, 50 mM Tris-HCl (pH 7.5), 5 mM EDTA, NP-40 (0.5% vol/vol), Triton X-100 (1.0% vol/vol)) on a shaker at room temperature for 60 min. Simultaneously, sheered chromatin samples (see above), blocking buffer and antibodies were added to a another polypropylene 96-well microplate (untreated) and incubated in ultrasonic bath for 60 min at 4 °C. The blocking buffer was aspirated from the protein A-coated plate, and the chromatin □+□ antibody mix from the untreated plate was transferred to the protein A-coated wells and incubated in the ultrasonic bath for 60 min at 4 °C. The wells were then washed 3 times with immunoprecipitation buffer followed by 3 washes with TE buffer. Finally, elution buffer containing 25 mM Tris base, 1 mM EDTA (pH10) with proteinase K 200 μg/ml was added to the wells, nutated for 30 s at 1400 rpms and incubated for 45 min at 55 °C and then 10 min at 95 °C. The 96-well plates were then briefly agitated and centrifuged for 3 min at ~□500 g at 4 °C and were used for PCR. The antibodies used for Matrix ChIP were: mCherry (Abcam ab167453), H3K27me3 (Active motif 39155), EZH2 (Cell Signaling D2C9), H3K27ac (Active motif 39133), p300(Santa Cruz sc-48343), Pol II 4H8 (Santa Cruz, sc-47701), Pol II Ser 5P+ (Santa Cruz), H3K4me3(Active motif 39159), Cas9 (Active motif 61757). Matrix ChIP experiments were performed in triplicate followed by qPCR in 4-8 replicates. ChIP-qPCR analysis and calculation are described below.

### ChIP-qPCR Calculations

All PCR reactions were run in quadruplicated using syber green. PCR calibration curves were generated for each primer set from a dilution series of total human genomic DNA. For each qPCR run the primer efficiency curve was fit to cycle threshold (CT) versus log (genomic DNA concentration) using an r-squared best fit. DNA concentration values for each ChIP and input chromatin DNA sample were calculated from their respective average Ct values as previously described^47, 84^. Final results are expressed as fraction of input DNA versus the normalization antibody H3 relative to no guide (-g).

### Cut and Run analysis

Cut and Run method and analysis was performed as previously described^65^. 1million WTC EBdCas9 cells gRNA transfected or not were harvested by centrifugation (600 g, 3 min in a swinging bucket rotor) and washed in ice cold phosphate-buffered saline (PBS). Nuclei were isolated by hypotonic lysis in 1 ml NE1 (20 mM HEPES-KOH pH 7.9; 10 mM KCl; 1 mM MgCl_2_; 0.1% Triton X-100; 20% Glycerol) for 5 min on ice followed by centrifugation as above. Nuclei were briefly washed in 1.5 ml Buffer 1 (20 mM HEPES pH 7.5; 150 mM NaCl; 2 mM EDTA; 0.5 mM Spermidine; 0.1% BSA) and then washed in 1.5 ml Buffer 2 (20 mM HEPES pH 7.5; 150 mM NaCl; 0.5 mM Spermidine; 0.1% BSA). Nuclei were resuspended in 500 μl Buffer 2 and 10 μl antibody was added and incubated at 4°C for 2 hr. Nuclei were washed 3 x in 1 ml Buffer 2 to remove unbound antibody. Nuclei were resupended in 300 μl Buffer 2 and 5 μl pA-MN added and incubated at 4°C for 1 hr. Nuclei were washed 3 x in 0.5 ml Buffer 2 to remove unbound pA-MN. Tubes were placed in a metal block in ice-water and quickly mixed with 100 mM CaCl_2_ to a final concentration of 2 mM. The reaction was quenched by the addition of EDTA and EGTA to a final concentration of 10 mM and 20 mM respectively and 1 ng of mononucleosome-sized DNA fragments from Drosophila DNA added as a spike-in. Cleaved fragments were liberated into the supernatant by incubating the nuclei at 4°C for 1 hr, and nuclei were pelleted by centrifugation as above. DNA fragments were extracted from the supernatant and used for the construction of sequencing libraries. We have also adapted this protocol for use with magnetic beads ^65^.

### RNA-seq data analysis

RNA-seq samples were aligned to hg19 using Tophat (version 2.0.13)^85^. Gene-level read counts were quantified using htseq-count using Ensembl GRCh37 gene annotations. Processed single cell RNA-seq data from Nakamura et al ^73^ were used. Only genes expressed above 10 Reads Per Million in 3 or more samples were kept. t-SNE was performed with the *Rtsne* package, using genes with the top 20% variance across samples. Cluster labels from Nakamura et al were used. A Principle Component Analysis (PCA) was performed using all of the cynomolgus monkey samples from Nakamura et al^73^ using R software. Genes used in the analysis were restricted to defined homologs expressed at non-zero Transcripts Per Million (TPM) in human *in vitro* cell lines, and in the preprocessed mouse and cynomolgus monkey single cell samples from Nakamura et al. RNA-seq data from human cell lines were corrected for batch effects using *ComBat*^86^. Human bulk RNA-seq samples were projected onto the PCA coordinate via matrix multiplication. Human, cynomolgus monkey and mouse RNA-seq data were separately centered and scaled within each species before PCA and projection was performed.

### Cell count

Cells were lifted using Versene and resuspended in media for cell count using hemocytometer. 100ul of cell suspension were applied to hemocytometer using 10X objective. 4 most outward corners were counted, averaged, multiplied by 10^4^, and multiplied by the dilution factor to get the final number of cells.

### Cell viability

Cell viability assay was measured by AlamarBlue (thermofisher scientific, DAL1025). Cells were induced and transfected with gRNA as discussed above. 6h after last transfection, 10X of viability reagent was added to media. Time “0” was measured 4h after. Fluorescence measurements were read on Perkin-Elmer Envision plate reader after transferring 50ul of media from each condition to 96 well microplate reader.

### CRISPR-based deletion of TBX18 TATA box region

For WTC EBdCas9 *TBX18* TATA box deletion (40 bp deletion), we used CRISPR/Cas9 to delete an 88bp region by using 2 gRNAs flanking TATA box 5’ and 3’ ends. Since gRNA 2 overlaps with *TBX18* g6 locus we reconstructed guide6 region using a ssDNA donor (168bp, IDT) that lacks the 40bp region around the TATA box (TATATGAC) but contains intact guide6 region and mutated PAM region. Two guides RNAs (Synthego) that are flanking TATA box region were designed using Benchling (Sup Table 8). Briefly, 0.5μM of spCas9 (sigma) were mixed with 1μM of gRNAs and incubated at RT for 15min. 1×10^6^ EBdCas9 cells were transfected with the above RNP complex and 100μM of ssDNA (HDR)(Sup Table 8) using AMAXA electroporation kit (LONZA) and plated in TeSR media (supplemented with 5μM ROCKi). Several EBdCas9 colonies were picked, followed by chromosomal DNA isolation (Quick Extract /Lucigen), PCR amplification (Sup Table 8) and PCR purification (ExoSAP-IT /Thermofisher). PCR products were sequenced to identify potential 40bp TATA box region deletions and g6 reconstruction. To produce a homozygous deletion of *TBX18* TATA box and guide 6 reconstruction a second hit of CRISPR/Cas9 was introduced using the same exact 2 guide RNAs and ssDNA as describe above. Genomic DNA sequencing analysis of selected clones confirmed homozygous deletion of 40bp of TATA box region on *TBX18*. To validate that the lines analyzed were homozygous TATA box deletion and guide6 reconstruction, we ligated PCR amplicons to pGEM-T (Promega) vector system and confirmed that all picked clones encompassed TATA box deletion and guide6 reconstruction using Sanger sequencing.

### ElemeNT Navigation Tool

TATA box prediction and other detection of core promoter elements were assayed using ElemeNT Navigation Tool^52^. Each analyzed sequence included 1000bp region. http://lifefaculty.biu.ac.il/gershon-tamar/index.php/element-description/element

### FANTOM5 CAGE Tool

Functional annotation of the mammalian genome was used to evaluate TBX18 transcription start site (TSS) against human RNAseq datasets. https://fantom.gsc.riken.jp/5/sstar/EntrezGene:9096

### QGRS Mapper Tool

Distribution of putative **Q**uadruplex forming **G**-**R**ich **S**equences (**QGRS**) in nucleotide sequences was predicted using QGRS mapper too. http://bioinformatics.ramapo.edu/QGRS/analyze.php

### Transcription factor prediction Tool

Prediction of transcription factor site scan on promoter region was analyzed using http://www.ifti.org/cgi-bin/ifti/Tfsitescan.pl

### Data availability

RNAseq dataset generated for this study is available from the NCBI GEO dataset under accession number: GSE155022. This Dataset corresponding to Figs: 5F, 5K, and Sup 5D.

## Supplemental Figure Legends

**Supplemental Figure 1:**
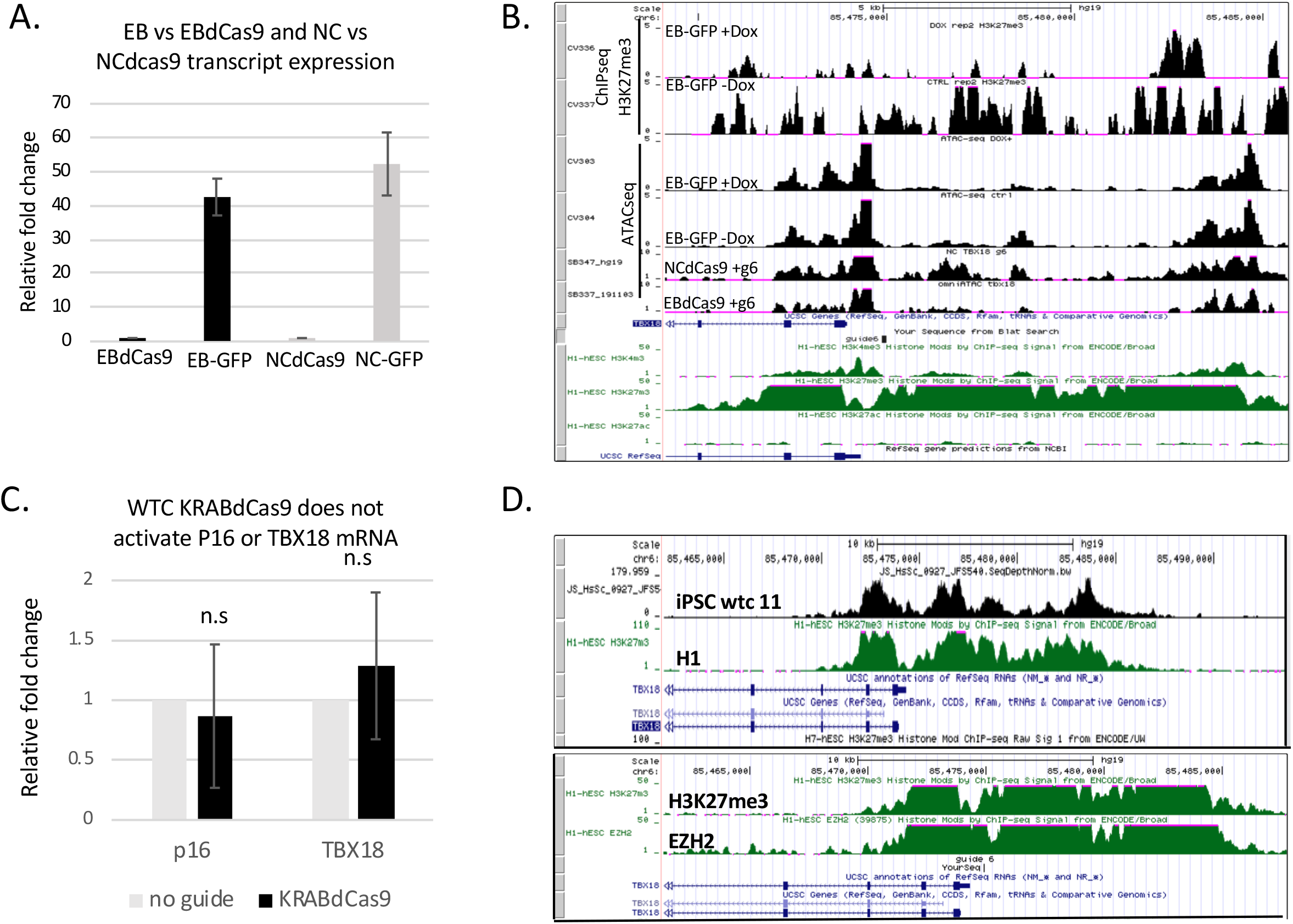
EBdCas9 upregulates TBX18 gene expression. **A**. EB-GFP and NC-GFP mRNA expression is x50 higher than EBdCas9 and NCdCas9 respectively. EB-GFP, NC-GFP, EBdCas9, and NCdCas9 were all induced with Dox for 3D and harvest for EB or NC transcript expression and measured using RT-qPCR. Beta-actin was used for normalization and all samples were compared to their no Dox cell lines counterparts. **B**. TBX18 has an open chromatin region. Genome browser view of EB-GFP+Dox and EB-GFP-Dox H3K27me3 ChIPseq analysis ^35^ and EB-GFP+Dox, EB-GFP-Dox, NCdCas9/g6, and EBdCas9/g6 ATACseq analysis. **C**. WTC KRABdCas9 does not activate TBX18. RT-qPCR analysis of TBX18 or p16 expression for KRABdCas9 normalized to beta-Actin and calculated as relative fold change compared to no guide (induced with Dox) of the respected cell line **D**. Top panel-iPSC WTC 11 EBdCas9 cell lines exhibits similar H3K27me3 tracks for TBX18 promoter region as H1 human embryonic stem cell line found in UCSC genome bowser. Bottom panel-EZH2 occupancy is similar to H3K27me3 in H1 hESC.

**Supplemental Figure 2:**
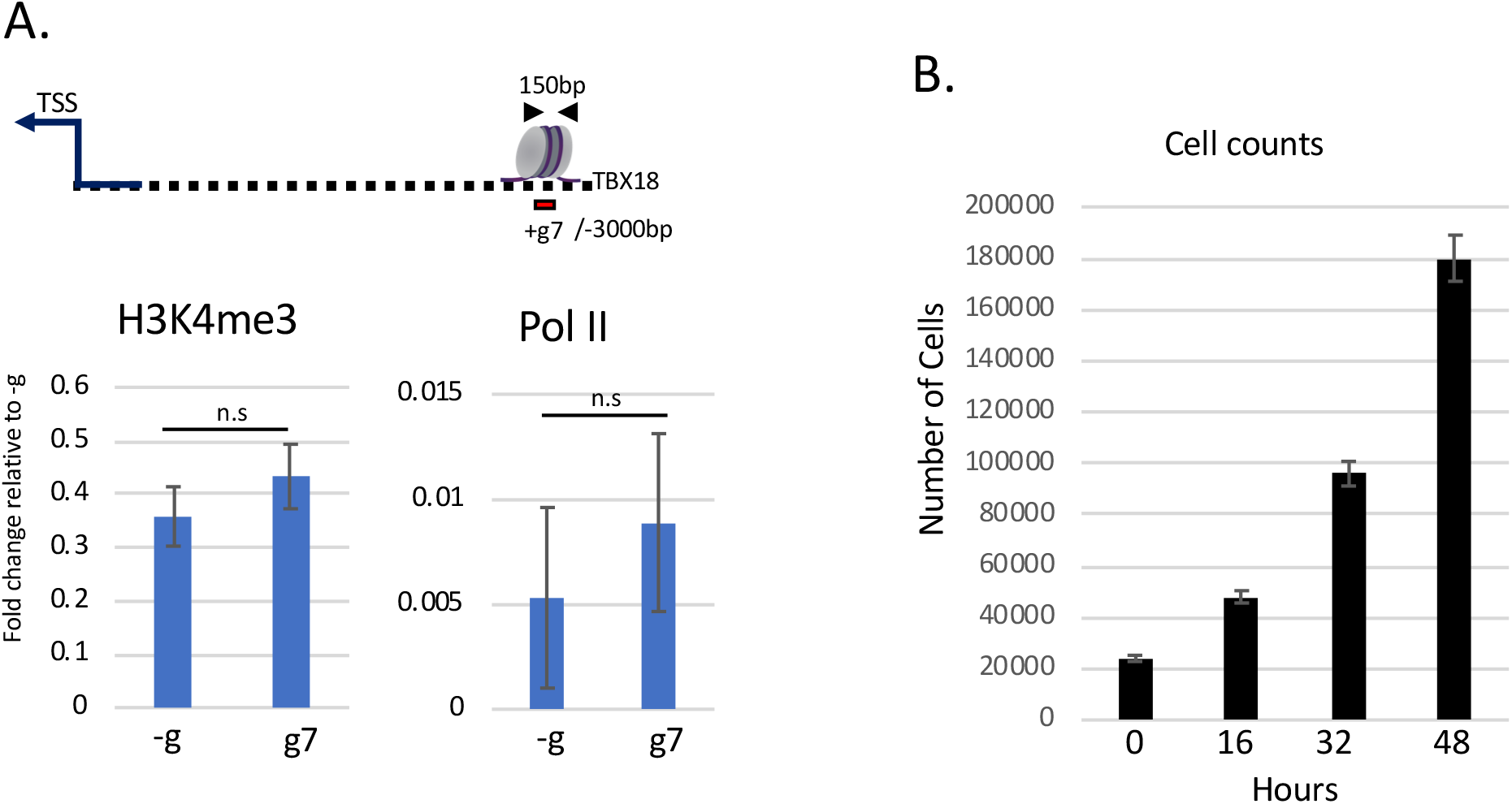
No epigenetic remodeling for EBdCas9/ TBX18 g7. **A.** ChIP-qPCR of TBX18 relative fold increase after 3D EBdCas9 dox induction and g7 RNA transfection (TBX18) normalized to fractioned Input/H3. Antibodies that were used for ChIP are listed above the graphs (H3K4me3, and Pol II) and the genomic region analyzed by qPCR includes TBX18 g7 flanking region locus. n=3 biological replicates. **B**. EBdCas9 doubling time is 16h. Cells were counted over a period of 48h and measured using nucleocounter.

**Supplemental Figure 3:**
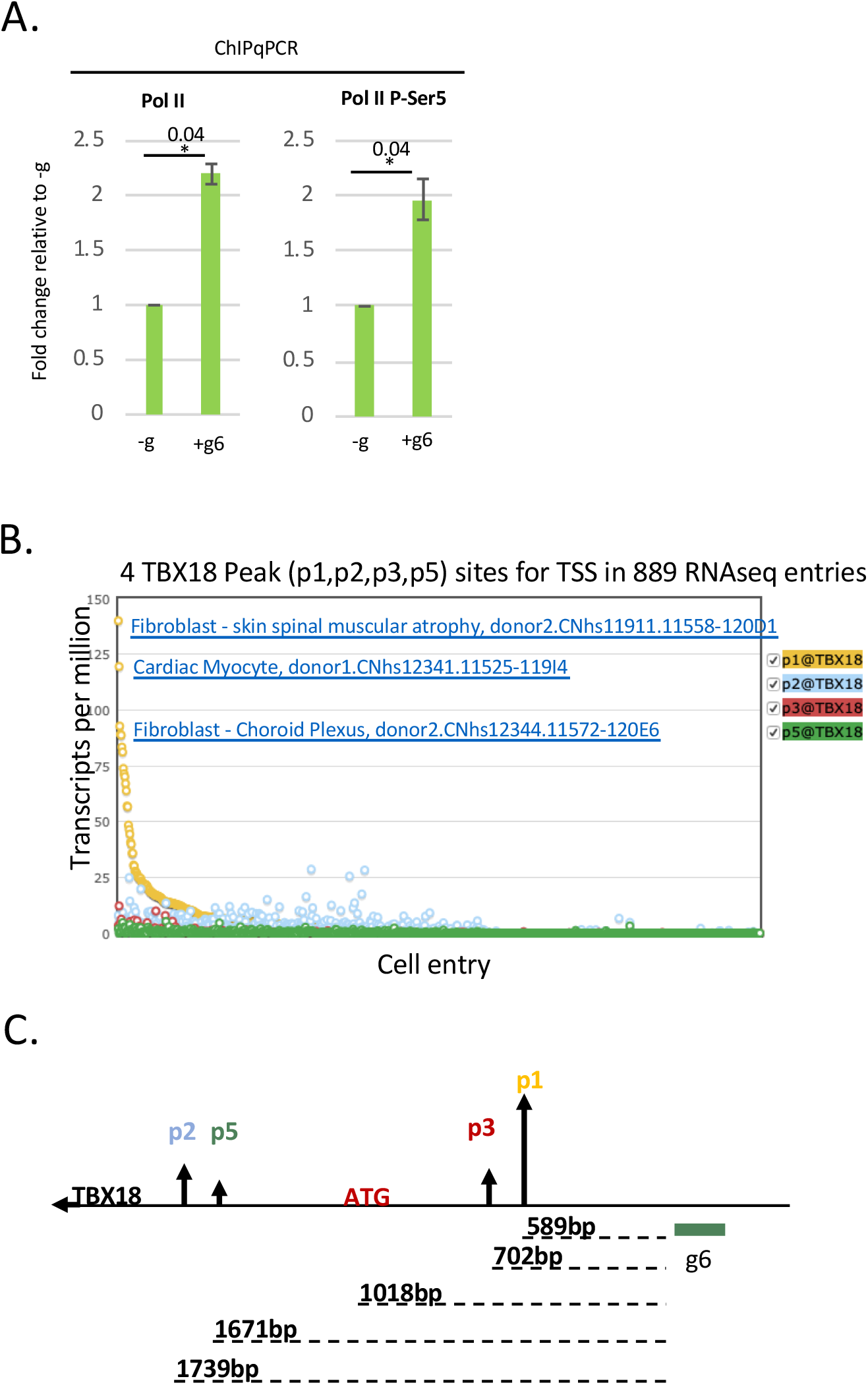
TBX18 TSS is predominantly 600bp downstream of guide 6. **A**.ChIP-qPCR of 3D induced EBdCas9 with TBX18 g6 RNA (+g6) transfected or not (-g). Normalized fractioned to Input/H3 and compared to no guide (-g). Antibodies that used for ChIP are listed above the graphs (Pol II CTD and Pol II P-Ser5) and the genomic region analyzed by qPCR flanking TBX18 g6 locus. \**p*<0.05, ** *p*<0.01, \*\*\**p*<0.001 one-way ANOVA performed. n=3 biological replicates. **B**. TBX18 input in FANTOM5-CAGE reveals 4 TSS peaks (p1, p2, p3, p5) measured by transcript expression (transcripts per million) among 889 human RNAseq datasets. The 3 top expressed datasets are displayed on the graph. **C**. Depiction of TBX18 4 TSS peaks with respect to guide6 region.

**Supplemental Figure 4:**
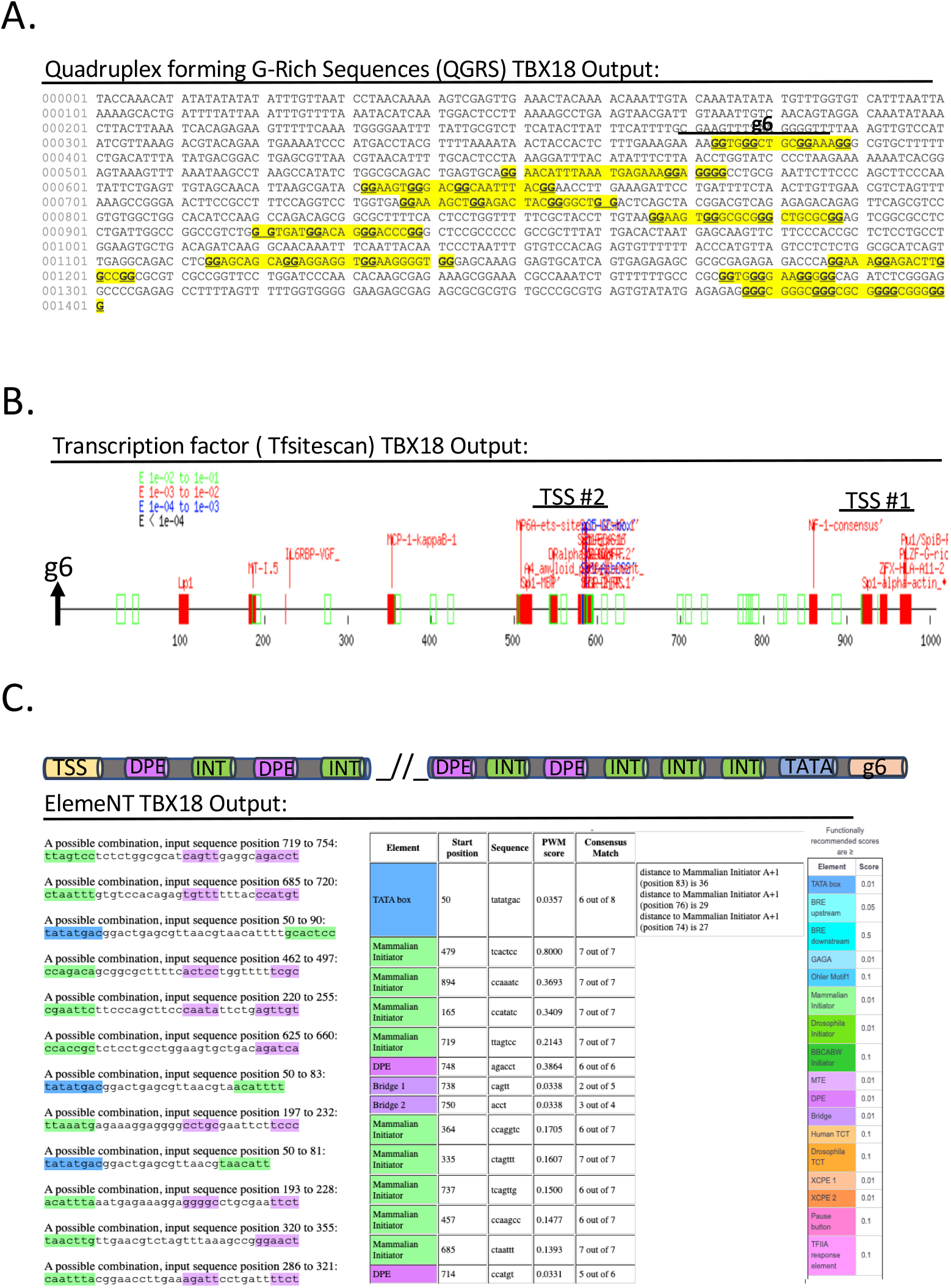
TBX18 promoter region is decorated with transcription elements. **A**. Snapshot of TBX18 1.0kb promoter region using the QGRS mapper tool. Highlighted in yellow are the G-quadruplex elements in 5’ to 3’ direction. **B**. Snapshot of TBX18 (1.0kb) promoter region using transcription factor site scan using Tfsitescan tool. g6, TSS#1 and TSS#2 are labeled on the axis. **C**. Snapshot of TBX18 TATA box and other transcriptional prediction elements using ElemeNT software tool.

**Supplemental Figure 5.**
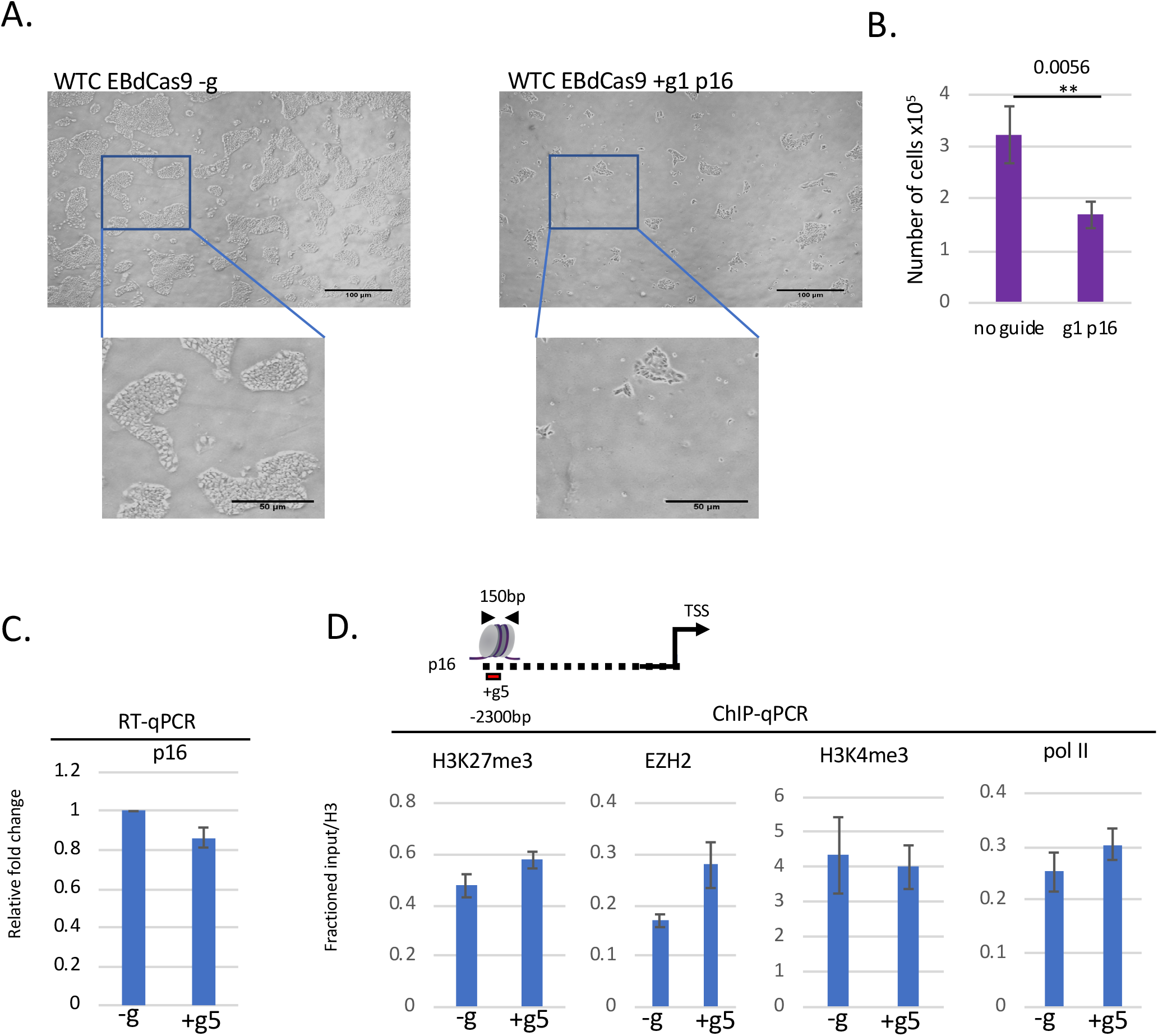
EBdCas9 upregulates p16 expression. **A-B.** EBdCas9/p16 g1 transfection results in cell proliferation reduction. (**A**) EBdCas9 brightfield after 3D p16 g1 transfection or no transfection (-g). Scale bar is 50*μ*m. (**B**) Total cell count of EBdCas9 after 3D p16 g1 transfection or no transfection (-g) in 9cm^2^ area field. n=3 biological replicates **C-D**. No epigenetic remodeling after EBdCas9 p16/g5 induction. **(C**) RT-qPCR or (**D**) ChIP-qPCR of p16 relative fold increase after 3D EBdCas9 dox induction and g5 RNA transfection (p16). RT-qPCR p16 normalized to beta-Actin and compared to no guide. ChIP-qPCR primer set target p16 g5 region (−2300bp) relative to TSS. Normalized to input and H3 and compared to −g relative fold change. Antibodies that were used for ChIP are listed above the graphs (EZH2, H3K27me3, H3K4me3, and Pol II) and the genomic region analyzed by qPCR includes p16 g5 locus. n=3 biological replicates.

**Supplemental Figure 6.**
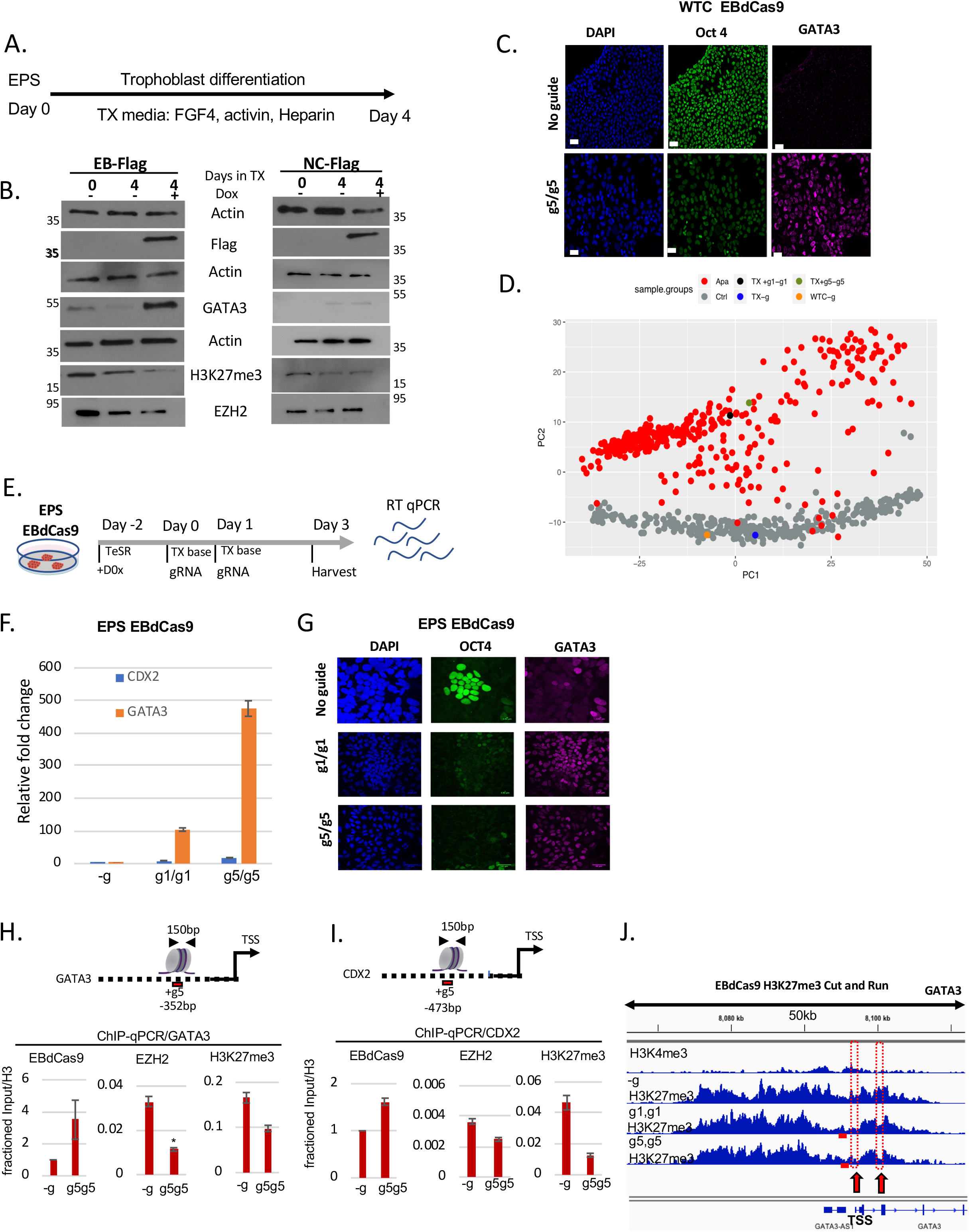
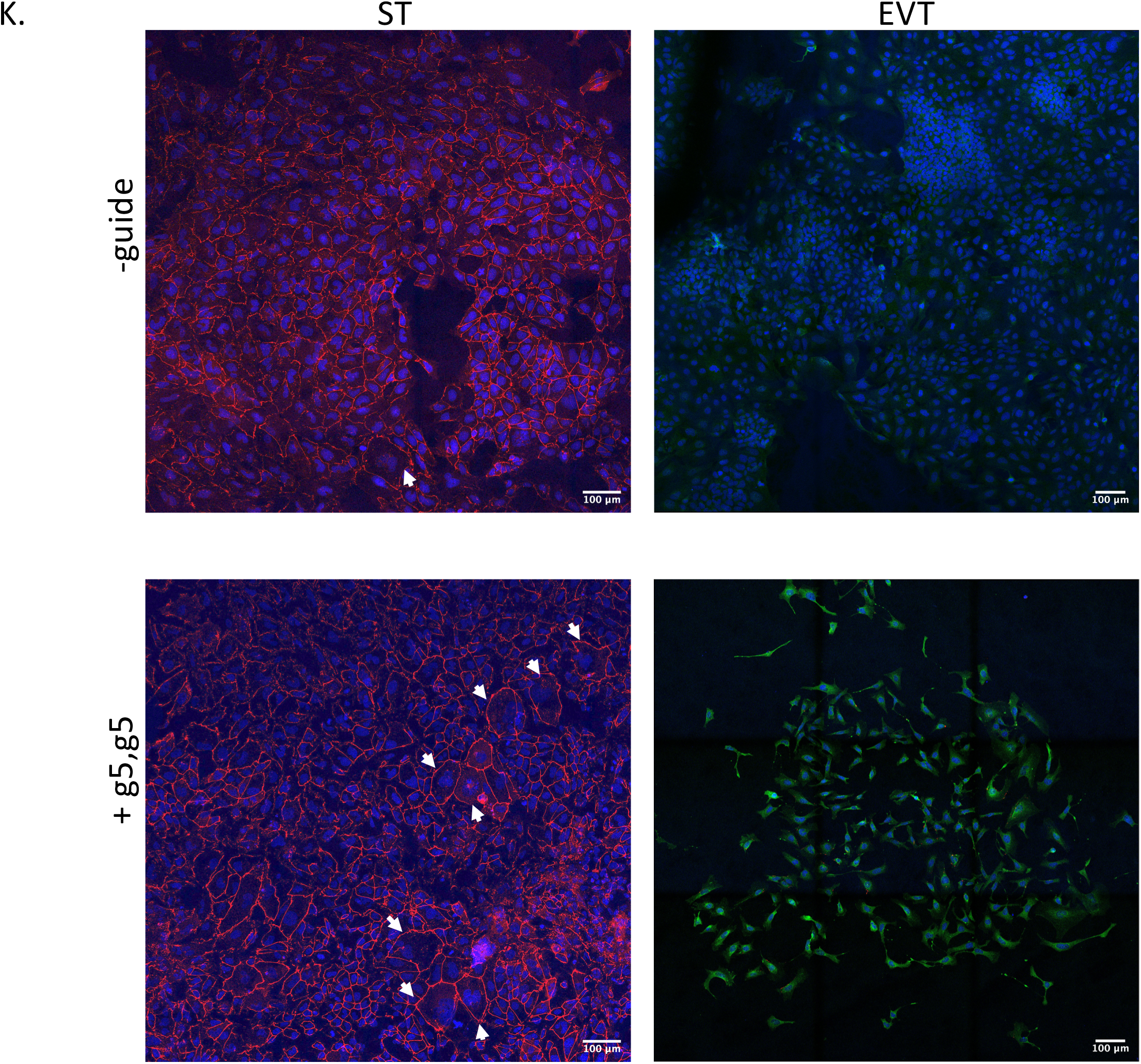
EBdCas9 *CDX2/GATA3* promotes trophoblast trans-differentiation. **A**. EPS EB-Flag and NC-Flag trophoblast differentiation time line. **B**. Immunoblot of EPS EB-Flag and NC-Flag with or without Dox after 4D of trophoblast differentiation in TX media or at EPS stage. **C**. Immunofluorescent imaging of 3D WTC EBdCas9 with (+) or without (-g) CDX2 and GATA3 g1g1 or g5g5 co-transfection. Blue-Dapi, Green-Oct4, Far red- GATA3; scale bar is 50*μ*m. **D**. PCA analysis of EBdCas9 CDX2/GATA3 gRNA co-transfection RNAseq compared to WTC dataset ^74^ **E**. WTC EBdCas9 cell line was reprogrammed to EPS, induced with Dox for 2D in TeSR and then switch to TX media (+Dox, no factors) while transfected with gRNA and harvest at 3D. **F**. RT-qPCR analysis of EBdCas9 EPS cells after co-transfection with g1/g1 CDX2/GATA3 or g5/g5 CDX2/GATA3 compared to no guide (-g) transfection. Normalized to beta-Actin. **G**. Immunofluorescent imaging of 3D EPS EBdCas9 with (+) or without (-g) CDX2 and GATA3 g1g1 or g5g5 co-transfection. Blue-Dapi, Green-Oct4, Far red- GATA3; scale bar is 50*μ*m. **H-I**. ChIP-qPCR analysis of 3D EBdCas9 co-transfected with g5/g5 CDX2/GATA3. Normalized to Input/ H3 and compared to −g relative fold change. Antibodies that were used for ChIP are listed above the graphs (Cas9, EZH2, H3K27me3) and the genomic region analyzed by ChIP-qPCR includes GATA3 g5 (**H**) and CDX2 g5 (**I**) loci. **J**. Cut and Run H3K27me3 analysis of 3D EBdCas9 co-transfected with g5/g5 CDX2/GATA3 or g1/g1 CDX2/GATA3. H3K27me3 tracks are normalized to IgG and displayed on Integrated Genome Viewer (IGV); 50kb GATA3 window viewer. Red dots demonstrate GATA3 TSS and exon 3 regions. **K**. Immunofluorescence of EBdCas9 3D post CDX2/GATA3 g5/g5 RNA transfection compared to no guide, and further 6 days (6D) differentiation to either EVT using 7.5mM TGFbi and 100ng/ml (NRG1) or ST using 2mM forskolin. Dapi- blue,ZO-1-red and (chorionic gonadotropin beta) CGB-green. Scale bar is 100*μ*m.

